# Alternative strategies of orthographic processing: the case of skilled deaf readers

**DOI:** 10.64898/2026.02.02.703016

**Authors:** Sendy Caffarra, Brendan Costello, Noemi Fariña, Jon Andoni Duñabeitia, Manuel Carreiras

**Affiliations:** University of Modena and Reggio Emilia, Department of Biomedical, Metabolic and Neural Sciences, 41125 Modena, Italy; Basque Center on Cognition, Brain, and Language, 20009 San Sebastian, Spain; Ikerbasque (Basque Science Foundation), Bilbao, Spain; Departamento de Biosanitaria, Facultad de Ciencias de la Salud, Universidad Internacional de La Rioja, Logroño, Spain; Centro de Investigación Nebrija en Cognición (CINC), Universidad Nebrija, Madrid, Spain; Departamento de Lengua Vasca y Comunicación, UPV/EHU, Bilbao, Spain

**Keywords:** reading, bimodal bilingualism, electrophysiology, orthographic processing, deaf readers

## Abstract

The cognitive factors that enable us to be proficient readers can greatly vary across individuals. The case of skilled deaf readers is emblematic as it shows that high reading performances can be achieved even when lifelong acoustic experience is absent or minimal. Here we present a set of experiments investigating how alternative strategies of orthographic processing can lead to high levels of reading proficiency. Four EEG studies compared behavioral and brain correlates of orthographic processing in skilled deaf readers and matched hearing controls. Using single word recognition and priming paradigms, we investigated two pillars of orthographic processing: letter identity and letter position. Our findings show that, although both groups had similarly accurate reading performance, skilled deaf readers were faster, and they consistently differ from hearing controls in the way they process letter identity. This group difference was observed in both lexical and sublexical tasks and was specifically related to the identity of orthographic representations, regardless of the visual form of the written stimuli (such as character visual similarity and letter case). These findings uncover alternative strategies that make possible high reading performance, even in the absence of acoustic experience.

**Public Significance Statement:** This research identifies alternative orthographic strategies that improve single-word reading efficiency and can potentially serve as effective compensatory tools when phonological processing is impaired.

## Introduction

Reading requires a complex set of cognitive skills that are subject to individual variability (Bonte & Brem, 2024; Stanovich, 1985). There is not a unique way to be a proficient reader and critical factors supporting efficient orthographic analysis can vary depending on people’s experience (Freed et al., 2017; Gordon et al., 2020). In this sense, deafness provides an extreme example of alternative cognitive strategies that can lead to proficient reading. Having a reduced or absent acoustic input is often associated with lower levels of reading proficiency as compared to hearing controls (Conrad, 1979; Qi & Mitchell, 2012), with a proficiency gap reported both in reading acquisition and in final attainment (Antia et al., 2020; Silvestri & Ehrenberg, 2021; Trezek & Mayer, 2019). Interestingly, despite this general trend, about 10% of deaf individuals are able to reach high levels of reading proficiency (Moreno-Pérez et al., 2015). This research work explores how these readers can achieve high levels of reading proficiency even when acoustic experience is low or absent. We ran a series of EEG studies to investigate the processing of visual word and pseudoword recognition in skilled deaf readers and tested how their reading mechanisms differ from those of matched hearing readers.

Previous studies on written language processing have pointed to phonological coding as the main difference between deaf and hearing readers. Compared to hearing controls, deaf readers show a reduced or even absent reliance on phonology during visual word recognition (Bélanger et al., 2012; Costello et al., 2021; Emmorey et al., 2013; Emmorey & Lee, 2021; Fariña et al., 2017; Holcomb et al., 2024; Lee et al., 2022; Ormel et al., 2010; Peleg et al., 2020; Rowley et al., 2025). Orthographic coding has been less investigated and the research evidence on this point is inconclusive. While some studies suggest that orthographic information is represented and processed similarly in hearing and skilled deaf readers (Fariña et al., 2017; Meade et al., 2019, 2020), another group of studies have reported an advantage for skilled deaf readers which could be related to 1) a faster or more efficient orthographic route (Bélanger & Rayner, 2015); 2) a greater reliance on orthographic cues or on orthographic-to-semantic mappings (Emmorey et al., 2016; Gutierrez-Sigut et al., 2019; Holcomb et al., 2024; Lee et al., 2022; Peleg et al., 2020; Winsler et al., 2023). Critically, none of the previous studies have extensively investigated different orthographic dimensions within both lexical and sub-lexical domains in the same group of deaf and hearing readers.

Here we present a series of four studies using electroencephalography (EEG) and behavioral scores that compared the time course of orthographic processing on the same groups of deaf skilled readers and matched hearing controls. In this set of experiments two main pillars of orthographic processing were investigated within the lexical and sublexical domains: letter identity and position. Our EEG findings consistently showed that skilled deaf readers differed from hearing controls in the way they processed letter identity in the early stages of lexical and sub-lexical analysis. Critically, although the time course of their orthographic analysis was different, both groups attained similar high levels of reading accuracy. This highlights the flexibility of the reading system in finding alternative orthographic strategies that can equally guarantee efficient reading.

### Orthographic processing in deaf and hearing readers

In the current study the processing of letter position and identity in deaf and hearing readers was tested using different paradigms (i.e., single word recognition, masked priming) and tasks (i.e., lexical decision, visual same-different matching). The role of letter position was assessed through transposed-letter (TL) items, which are created by transposing two letters of an original target word (e.g., TL: tefélono; original word: teléfono, *phone* in Spanish). Letter identity was also manipulated in visual word recognition through replaced-letter (RL) items, which are created by replacing two letters of an original target word (e.g., RL: tehékono; original word: teléfono, *phone* in Spanish). As a result of the transposition or the replacement, lexical access becomes more difficult, with RL and TL showing lower accuracy and/or slower reaction times in lexical decision tasks as compared to matched control words and/or “normal” nonwords (Andrews, 1996; Chambers, 1979; Grainger et al., 2012; Lupker & Pexman, 2010). Also, TL are usually more difficult than RL as they more likely activate the corresponding existing orthographic representation which contains the same letters in a different order (e.g., as compared to the RL tehékono, the TL tefélono will more easily activate the real Spanish word teléfono; Andrews, 1996; C. E. Lee et al., 2024; Mirault & Grainger, 2021). Behavioral studies comparing matched skilled deaf readers and hearing controls showed that deaf readers can achieve similar (and sometimes even higher) behavioral performance to hearing controls in a lexical decision and priming paradigms involving TL, RL and real words (Costello et al., 2021; Fariña et al., 2017; Gutierrez-Sigut et al., 2022; Lee et al., 2022; Meade et al., 2020; Peleg et al., 2020). This suggests that when the outcome of the orthographic processing is behaviorally measured, the relative contribution of letter identity and letter position in text recognition might not differ between skilled deaf readers and hearing controls. However, even if the performance is similarly accurate between groups, there is still the possibility that the cognitive processes supporting these behavioral performances are different. To address this point, high temporal resolution techniques, such as EEG, are essential to examine the time course of orthographic processing of both groups. Event-related potentials (ERPs) are electrophysiological responses time-locked to the presentation of an external event (e.g., written word presentation) that provide information about how brain responses of orthographic analysis unfold over time, millisecond by millisecond. Three main ERP components have been reported to be sensitive to the effects of letter position and letter identity in priming paradigms: the N250, the N400 and the Late Positive Complex (LPC). The N250 is a negative component peaking around 250 ms after stimulus onset which is associated with the sub-lexical analysis of text (Grainger et al., 2006; Holcomb & Grainger, n.d., 2007). Its amplitude modulates based on orthographic properties of the stimuli (e.g. variations in letter position and identity), and it is usually smaller for pseudowords (i.e., TL and RL) as compared to words (Coch & Mitra, 2010; Zhang et al., 2021). In priming paradigms, target stimuli elicited smaller N250 when preceded by transposed-letter primes than by replaced-letters primes suggesting a stronger target pre-activation and a flexible representation of letter position (Carreiras et al., 2009; Duñabeitia et al., 2009; Grainger et al., 2006; Meade et al., 2021; Pegado et al., 2021; Perea & Lupker, 2004). The N400 represents a negative deflection peaking around 400 ms over centro-posterior sensors. Its amplitude modulates based on the lexical status of the target stimulus with greater amplitude for pseudowords as compared to words (Coch & Mitra, 2010; Kutas & Federmeier, 2011; Vergara-Martínez et al., 2013; Zhang et al., 2021). The LPC is a late positive deflection peaking around 600 ms after stimulus onset, which is posteriorly distributed. Its amplitude modulates based on the lexicality of the stimulus and it is usually greater for novel items, such as pseudowords, than known words (Segalowitz et al., 1997). This LPC effect reflects late controlled processes of repair, monitoring and re-processing that are decision-dependent (van de Meerendonk et al., 2013; Vissers et al., 2008; Yang et al., 2019).

EEG studies that have been focused on the role of letter position during orthographic coding did not report consistent ERP differences between skilled deaf readers and hearing controls. For instance, a lexical decision study showed that TL elicited greater negative responses (N250 and N400) as compared to matched words and this effect did not differ between deaf and hearing controls (Lee et al., 2022). Similarly, a study using a masked priming paradigm showed that, when target words were preceded by TL primes, they elicited smaller negative effects (N250 and N400) as compared to the RL, suggesting that TL more easily activated the corresponding orthographic representations facilitating the recognition of the upcoming target word (Meade, 2020). Critically, this effect did not differ between skilled deaf readers and hearing controls, indicating a similarly flexible representation of letter position (Meade, 2020). However, a recent masked priming study reported an early group difference with a reversed and delayed N250 priming effect for deaf readers as compared to hearing controls, possibly signaling a more robust reliance on whole-word orthographic representations (Holcomb et al., 2024).

On the other hand, consistent group differences have been reported in the processing of letter identity of skilled deaf and hearing readers. An ERP priming study on a lexical decision comparing words and RL found a lexicality effect on the N400 for both skilled deaf and hearing readers, but only deaf readers showed sensitivity to RL orthographic similarities with the original target words (Gutierrez-Sigut et al., 2022). Similarly, an ERP priming study compared target words preceded by RL primes that could be existing words (i.e., orthographic neighbors of the target) or pseudowords (Meade, 2019). In deaf and hearing controls, target words preceded by neighboring word primes elicited a greater N400 effect as compared to those preceded by pseudowords, suggesting an increased effort in the lexical access of the target probably due to lexical competition. Critically, the effect was more anteriorly distributed in hearing controls, indicating stronger lexical competition due to the activation of orthographic, as well as phonological representations.

Hence, these findings point to the possibility that the way letter identity is treated during word recognition changes depending on our acoustic experience. Here, we examine the time course of letter identity analysis in four different experimental designs to provide a more nuanced understanding of alternative orthographic strategies ensuring high levels of reading proficiency. These studies were not preregistered.

### Experiment 1

Experiment 1 examines the impact of letter identity and letter position during visual word recognition processing in deaf and hearing readers. We directly compared RL, TL and matched words in a single-word lexical decision paradigm. Based on previous literature, greater N400 effects (and LPC) were expected for pseudowords (RL and TL) compared to words (Coch & Mitra, 2010; Gutierrez-Sigut et al., 2022; Kutas & Federmeier, 2011; Lee et al., 2022; Vergara-Martínez et al., 2013; Zhang et al., 2021). We predicted that deaf and hearing readers would differ in the way they treat orthographic information over time, and these group differences should be more evident for analysis of letter identity (Gutierrez-Sigut et al., 2022; Lee et al., 2022).

#### Methods

##### Participants

Twenty severely (70-90 dB) to profoundly (>90 dB) deaf adults (14 females; age range: 23-45 years old; mean: 33; SD: 7) participated in the study. All participants were born deaf or lost their hearing before the age of 3; none had cochlear implants. They learned Spanish Sign Language before the age of 10 and used it as the main language for communication. Most of them learned to read at an early age, at school, except two individuals who learned to read after the age of 16. Twenty hearing Spanish readers (10 females; age range: 20-42 years old; mean: 29; SD: 6) were included as a control group. All deaf participants and hearing controls were skilled readers in Spanish with performances above the 75th centile of the ECL-2 Test (De la Cruz, 1999) and the two groups did not differ in non-verbal intelligence, Spanish reading comprehension, and Spanish vocabulary size (see Table 1). Before taking part in the experiment, all participants signed an informed consent form that was approved by the BCBL Ethics Committee (number of protocol: 7212A). The experiment was performed following the BCBL Ethics Committee’s guidelines and regulations.

**Table 1.**
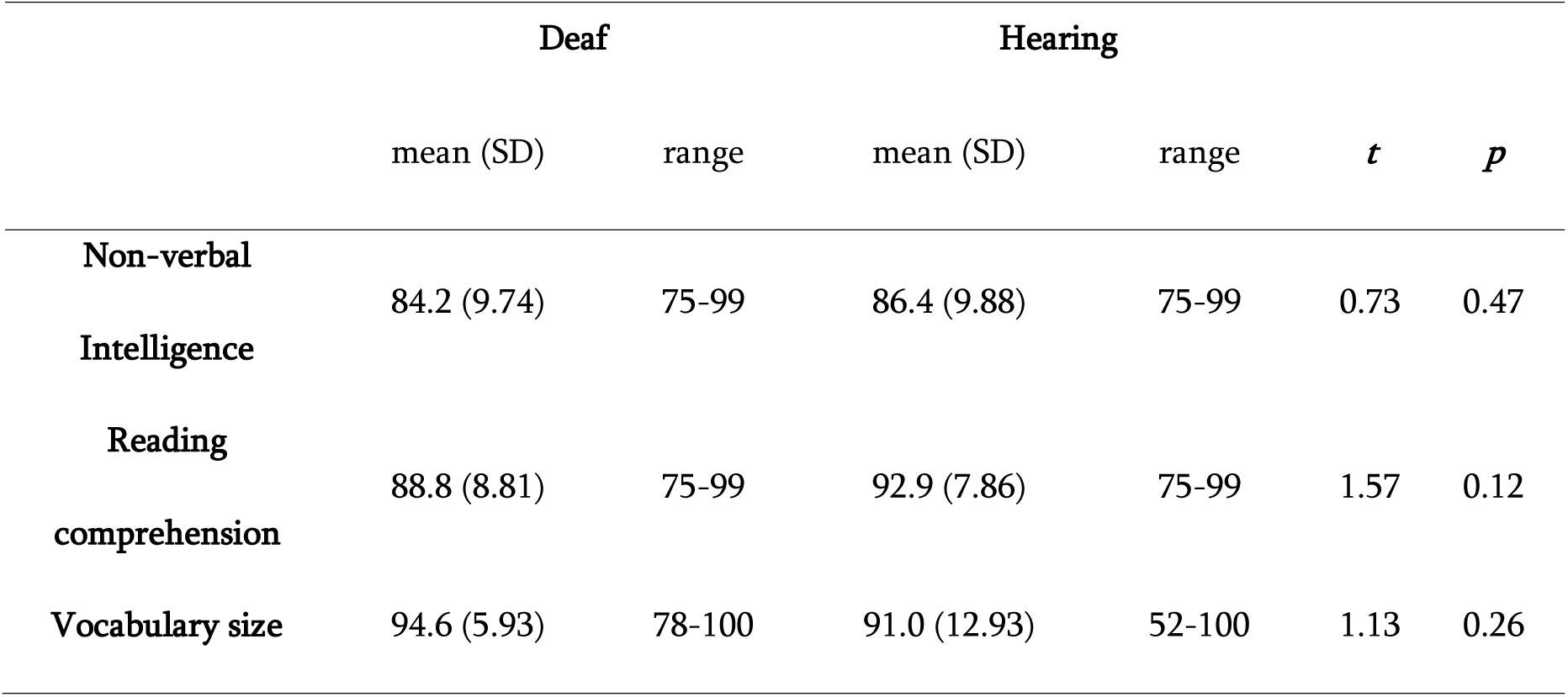
Characteristics of the experimental and control groups. Non-verbal intelligence was measured with the Raven Progressive Matrices test (Raven et al., 1998), where participants had to identify the missing item of a sequence of figures. Spanish reading comprehension was assessed through the ECL-2 (De la Cruz, 1999), a standardized reading test consisting of five short paragraphs followed by a total of 27 multiple-choice questions evaluating word meaning, synonyms, antonyms, sentence and text content. Vocabulary size was measured using the Spanish version of the LexTALE (Izura et al., 2014; Lemhöfer & Broersma, 2012), a lexical decision test consisting of 60 real words and 30 nonwords providing a good estimate of language knowledge (de Bruin et al., 2017). All scores are percentages. Results of two-tailed t-tests are reported for each group comparison.

##### Materials

For word trials, we selected 80 Spanish words between eight and ten letters long (mean number of letters 8.73; average log frequency 3.88, range 3.45–4.48) from the EsPal database (Duchon et al., 2013). For nonword trials, 80 Spanish base words between eight and ten letters long (mean number of letters 8.74) with a similar frequency to the first set were also selected (mean log word frequency 3.91, range 2.97–5.12). These words were then used to create two types of nonwords: (1) transposed-letter (TL) nonwords, in which the position of two non-adjacent consonants was swapped (e.g., *mecidina* from the base word *medicina* [medicine]); and (2) replaced-letter (RL) nonwords in which the two critical consonants were substituted by others with a similar visual shape as those used in the transposed-letter non-word (e.g., the pseudoword *mesifina* created from the same base word). Two lists were constructed such that each base word used to generate the nonwords was presented once in each list, either as a transposed-letter nonword or as a replace-letter nonword. Participants were randomly assigned to the two lists. In total, each participant completed 160 trials: 80 trials with words and 80 trials with nonwords (40 transposed-letter nonwords and 40 replaced-letter nonwords).

##### Procedure

The experiment was run in a silent and dimly lit room using Presentation software (version 0.70). Each trial began with a fixation cross at the centre of the screen for 500 ms, followed by a blank screen for 200 ms. Then, a stimulus (word or nonword) was presented in lowercase font (45-pt. Courier New, subtending a visual angle of 8.5 to 9.5 degrees) for 1500 ms or until the participant’s response (see Fig. 1A). Participants were instructed to press one of two buttons on the keyboard (‘M’ and ‘Z’) to indicate whether the letter string was a word or a nonword. They were asked to respond as quickly and accurately as possible. The order of presentation of the stimuli was randomized for each participant, and the two response buttons were counterbalanced for word and nonword responses. Participants completed eight practice trials before starting the experiment.

**Figure 1.**
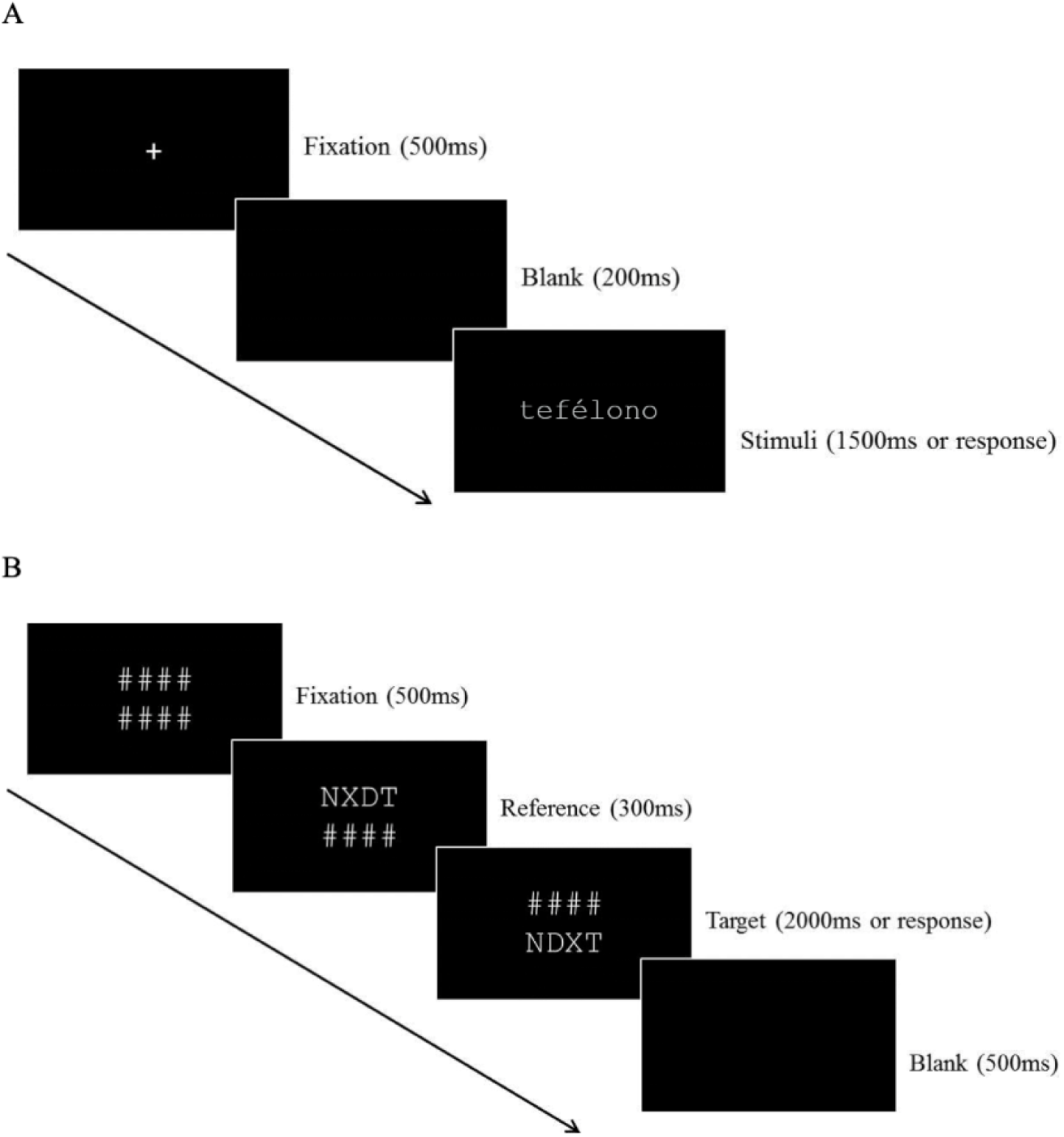
A: a trial sequence for the lexical decision task (Experiment 1). B: a trial sequence for the letter string matching task (Experiments 2-4).

##### EEG recording and analysis

The electroencephalogram (EEG) was recorded with a 32-channel Brain-Amp system (Brain Products GmbH) at a 500 Hz sampling rate. Scalp voltages were collected from 27 Ag/AgCl electrodes placed in an EasyCap recording cap (Fp1, Fp2, F7, F8, F3, F4, FC5, FC6, FC1, FC2, T7, T8, C3, C4, CP5, CP6, CP1, CP2, P3, P4, P7, P8, O1, O2, Fz, Cz, Pz). Two external electrodes were placed on the mastoids, with the right one as the online reference. An additional electrode at FCz served as ground, and four electrodes (two above and below the right eye, and two on the ocular canthi) recorded the electro-oculogram (EOG). Impedance was kept below 5KΩ for mastoids and scalp electrodes, and below 10KΩ for EOG electrodes. The EEG signal was re-referenced offline to the average activity of the two mastoids. A notch filter and a bandpass filter were applied (50 Hz, 0.01–30 Hz, 12 dB/octave). Ocular artifacts were corrected using Independent Component Analysis (ICA): the EEG of each subject was decomposed into independent components and those that explained the highest variance of EOG channels were identified and visually inspected before being removed from the original signal. ERPs time-locked to the onset of the stimulus presentation were obtained only for the trials followed by a correct behavioral response. EEG epochs of 1200 ms time-locked to the target presentation onset were obtained (including 200 ms pre-stimulus baseline). Residual artifacts exceeding ± 100 µV in amplitude were removed. The remaining clean epochs were averaged across conditions, with a baseline correction between – 200 and 0 ms. Statistical analyses were conducted on both behavioral and ERP data.

We analyzed the data using R and the lme4 package (Bates et al., 2015; R Core Team, 2013). For the behavioral data, we calculated error rates and reaction times (RTs). Trials with incorrect responses were excluded from RT analysis. RTs that were more than 2.5 SDs away from the mean of each condition and each participant were also excluded from the analysis of the response latencies. A by-subject repeated-measures ANOVA was conducted on the accuracy scores and the RTs including Group (Deaf, Hearing) as a between-subject factor, and Word Type (Word, TL, RL) as a within-subject factor. For the EEG data, a repeated-measures ANOVA was conducted on the average ERP amplitude within four time windows: 300-450 ms, 450-600 ms, 600-800 ms and 800-1000 ms. The average activity of three contiguous electrodes was calculated resulting in nine clusters: Left Anterior (F3, F7, FC5), Left Central (C3, T7, CP5), Left Posterior (P3, P7, O1), Medial Anterior (Fp1, Fp2, Fz), Medial Central (FC1, FC2, Cz), Medial Posterior (CP1, CP2, Pz), Right Anterior (F4, F8, FC6), Right Central (C4, T8, CP6), Right Posterior (P4, P8, O2). To probe the scalp distribution of any effects, these clusters were included in the statistical analyses as different levels of two topographical factors: Anteriority (Anterior, Central, Posterior) and Laterality (Left, Middle, Right). Each ANOVA included Group (Deaf, Hearing) as a between-subject factor and Word Type (Word, TL, RL) as a within-subject factor. The Greenhouse–Geisser procedure was applied when the sphericity assumption was violated. Effects of topographical factors will be reported only when they interact with the experimental factors. In both behavioral and EEG analyses, two-tailed t-tests were conducted as follow-up analyses and corrected for multiple comparisons (Hochberg, 1998). Linear mixed effects (LME) models for accuracy, RTs and ERPs led to similar results and are reported in full in the Supplementary materials. Preprocessed data, code and experimental materials are available here <u>osf.io/x3g5z</u>

#### Behavioral results

##### Accuracy

A Word Type effect was reported (Word Type: *F(2,76)*=82.502, *p*<.001), indicating that TL was less accurate than the other two conditions (TL vs words: *t(39)*=9.190, *p*<.001; TL vs RL: *t(39)*=9.229, *p*<.001; see Figure 2). No Group main effect or interaction was observed.

**Figure 2.**
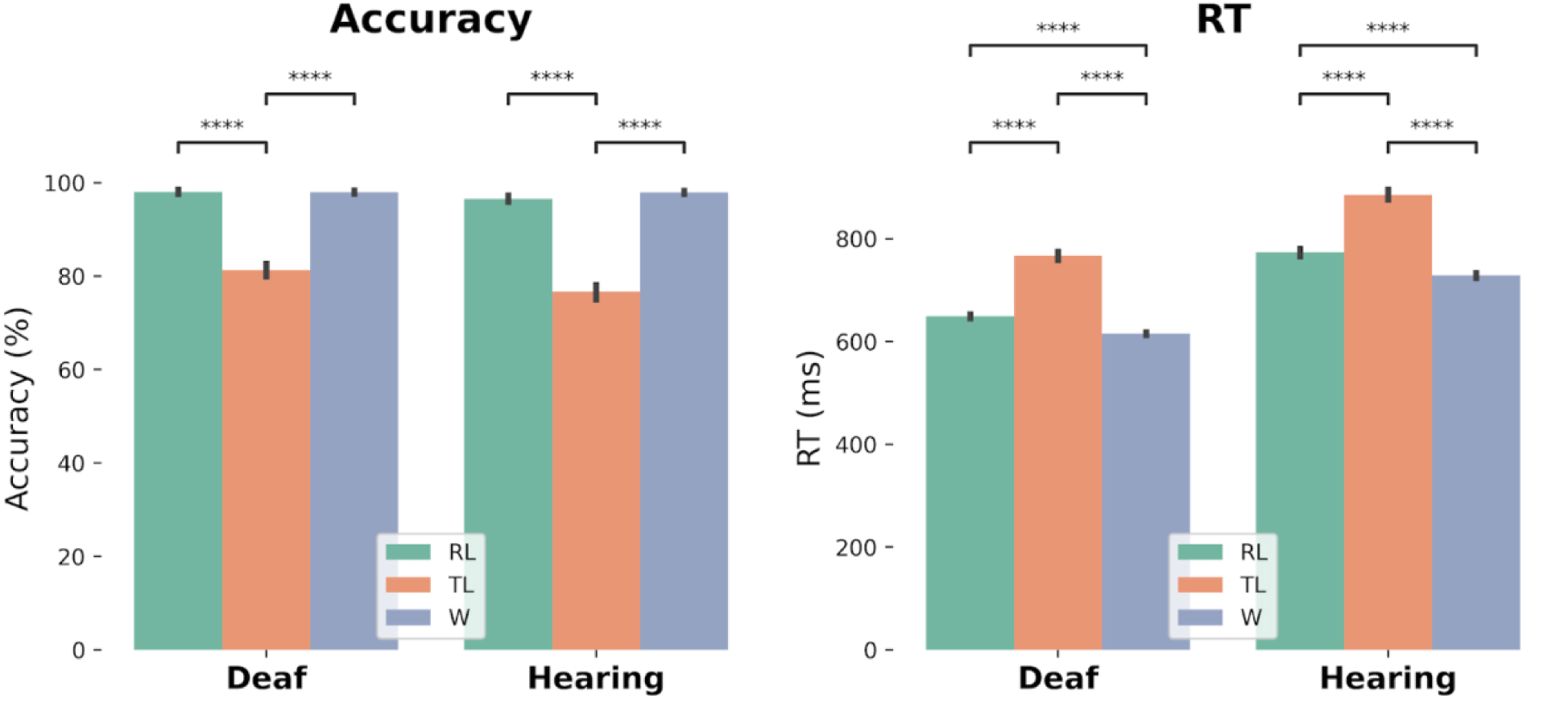
Average accuracy and reaction times for each Group and Word Type. Error bars represent +/- 1 SE. TL was the condition showing the lowest accuracy and the slowest responses. ****: p<0.0001.

##### RT

Deaf readers were overall faster than hearing controls (Group: *F(1,38)*=11.895, *p*=.001). The Word Type effect was also significant (Word Type: *F(2,76)*=79.479, *p*<.001), indicating that words were the fastest, followed by RL and then TL (words vs RL: *t(39)*=4.616, *p*<.001; RL vs TL: *t(39)*=8.970, *p*<.001; words vs TL: *t(39)*=10.071, *p*<.001; see Figure 2). The interaction Group x Word Type was not significant.

##### ERP results

No significant effects involving our experimental conditions were found before 300 ms.

###### 300-450 ms

The pseudowords conditions (RL and TL) elicited a greater negativity as compared to the word condition (Word Type: *F(2,76)*=13.928, *p*<.001), and this N400 effect was greater over central sites (Word Type x Anteriority: *F(4,152)*=4.880, *p*<.01; Word Type x Anteriority x Laterality: *F(8,304)*=2.798, *p*<.05; RL vs words reached its maximum over anterior-central medial sites, where *ts*>5, *ps*<.001; TL vs words reached its maximum over central medial sensors, with *ts*>3.5, *ps*<.01). No significant interaction between Word Type and Group was found in this time window.

###### 450-600 ms

Although an N400 effect was still present in this time window (Word Type: *F(2,76)*=29.663, *p*<.001; Word Type x Anteriority: *F(4,152)*=9.977, *p*<.001; Word Type x Laterality: *F(4,152)*=3.253, *p*<.05; Word Type x Anteriority x Laterality: *F(8,304)*=6.247, *p*<.001), it significantly varies as a function of group (Group x Word Type x Anteriority: *F(4,152)*=4.251, *p*<.05). Follow up analyses showed that while in deaf participants all pseudowords (RL and TL) were still eliciting a greater central-posterior N400 (TL vs words and RL vs words differed over central: *ts*>2, *ps*<.05 and posterior sensors: *ts*>3, *ps*<.01), in hearing controls only the TL condition showed a greater N400 effect as compared to the other two conditions (TL vs RL and TL vs words differed over anterior: *ts*>5, *ps*<.001, central: *ts*>4, *ps*<.01, and posterior sensors: *ts*>3, *ps*<.05). Hence, the two groups critically differed in the RL condition, which showed a longer-lasting N400 effect only in the case of deaf readers.

###### 600-800 ms

Again a Word Type effect was reported (Word Type x Anteriority: *F(4,152)*=7.781, *p*<.001; Word Type x Anteriority x Laterality: *F(8,304)*=2.868, *p*<.05) and it significantly interacted with the factor Group (Group x Word Type: *F(2,76)*=6.535, *p*<.01; Group x Word Type x Anteriority: *F(4,152)*=2.860, *p*<.05) suggesting that only in the deaf reader group did pseudowords (RL and TL) elicit a greater anterior-central positivity as compared to words (RL vs words and TL vs words showed a differences in the anterior: *ts*>3, *ps*<.05, and central sites: *ts*>3, *ps*<.05). However, this effect was not fully replicated in mixed linear models and will not be further discussed (see Supplementary Materials S2).

###### 800-1000 ms

Both groups showed a greater positivity for TL than words (Word Type: *F(2,76)*=19.639, *p*<.001; Word Type x Anteriority: *F(4,152)*=17.974, *p*<.001; Word Type x Anteriority x Laterality: *F(8,304)*=5.105, *p*<.001). However, the topographic distribution of the effect was more widespread for deaf readers as compared to hearing controls (Group x Word Type x Anteriority: *F(4,152)*=3.854, *p*<.05: TL vs words was significant on all scalp for deaf readers, *ts*>3, *ps*<.05, but only on posterior sensors in hearing controls, *t*>5, *p*<.001).

#### Discussion

Experiment 1 investigated the relative contribution of letter position and letter identity during a lexical decision task performed by skilled deaf readers and hearing controls. Behavioral data showed similar high levels of accuracy across groups, with TL being the most difficult condition for both deaf and hearing readers (i.e., the slowest and least accurate, as in Chambers, 1979; Fariña et al., 2017; O’Connor & Forster, 1981). In line with previous literature (Clark et al., 2016; Fariña et al., 2017; Morford et al., 2017, 2019; Villwock et al., 2021), deaf readers were faster than hearing controls, which has been related to greater sensitivity to visual-orthographic cues (Bélanger & Rayner, 2015; Emmorey et al., 2016; Gutierrez-Sigut et al., 2019; Lee et al., 2022; Meade et al., 2019; Morford et al., 2017), absence of phonological competition (Costello et al., 2021), and/or a visual attention benefit (Bélanger et al., 2017; Pavani & Bottari, 2011).

In the ERPs, both groups showed a N400 effect for pseudowords (TL and RL) as compared to words, suggesting that both types of pseudowords required increased cognitive resources during lexical access as compared to words (Coch & Mitra, 2010; Kutas & Federmeier, 2011; Vergara-Martínez et al., 2013; Zhang et al., 2021). Critically, deaf readers and hearing controls differed in the way they treated RL during lexical access (indexed by the N400 effect; Gutierrez-Sigut et al., 2022; Lee et al., 2022). As compared to hearing controls, deaf readers showed a longer-lasting effect of letter replacement, indicating a more sustained sensitivity to letter identity during lexical analysis (possibly related to lexical selection and stronger competition from orthographic neighbors; Yan et al., 2015).

In the later time window, both groups showed an LPC effect for TL as compared to the other two conditions (RL and words), which was in line with the behavioral results and indicated increased costs of repair and re-analysis (possibly due to letter swapping attempts with TL). Deaf readers showed a broader topographic distribution of this late LCP effect, suggesting a larger recruitment of neural sources as compared to hearing controls during late phases of re-analysis of letter position.

Overall, Experiment 1 showed that, within a lexical domain, deaf readers and hearing controls reached high levels of accuracy. However, ERP responses showed that they could do so by using different strategies of analysis of both letter identity and letter position. Experiment 2 tested whether these research findings could be generalized to the orthographic domain.

### Experiment 2

Experiment 2 tested whether skilled deaf readers and hearing controls adopted different strategies of orthographic processing even when written stimuli never corresponded to a lexical entry and the task did not require judging the lexical status of the target stimuli. To this aim, consonant strings were presented in a masked priming design where the target was a letter string that could be preceded by an identical prime, an RL version of the target or a TL version of the target. Behavioral and ERP data were collected to check for group differences in the time course of the orthographic analysis, as well as in the final response outcomes.

#### Participants

The participants were the same as those in Experiment 1.

#### Materials

The stimulus pairs in the perceptual matching task consisted of two character strings of four letters: a prime and a target. The uppercase consonants B, C, D, F, G, H, J, K, L, M, N, P, Q, R, S, T, V, W, X, Y and Z were used in the strings. The prime was either identical to the target (matched items) or differed from the target (mismatched items). Mismatched primes were created by modifying the internal characters of the target string, either by interchanging their position to create transposed-letter strings (TL, e.g., NDXT-NXDT), or by substituting in other characters to create replaced-letter strings (RL, e.g., NFLT-NXDT). No item repetition occurred within the experiment other than that each target string appeared twice (once with a matching prime and once with a mismatching prime). Two lists were constructed with an equal number of TL and RL primes, and such that across both lists, a given target had a mismatch with a TL prime in one list and with a RL prime in the other list. Participants were randomly assigned to one of the two lists. In total, each participant completed 160 trials: 80 trials with matched strings and 80 trials with mismatched strings (40 TL primes and 40 RL primes).

#### Procedure

The experiment was run in a silent and dimly lit room using Presentation software (version 0.70). Each trial began with the presentation of two lines of four hash marks above and below the center of the screen (see Fig. 1B) for 500 ms. Then, the prime (35-pt. Courier New, subtending a visual angle of 3.4 degrees) replaced the top row of hash marks for 300 ms while the bottom row remained visible. Finally, the prime was replaced by hash marks while the target string appeared on the bottom row for 2000 ms or until the participant responded. Participants were instructed to press one of two buttons on the keyboard (‘M’ and ‘Z’) to indicate whether the letter strings matched or mismatched. They were asked to respond as quickly and accurately as possible. The order of presentation of the stimuli was randomized for each participant, and the two response buttons were counterbalanced for match and mismatch responses. Participants completed eight practice trials before starting the experiment.

#### EEG recording and analysis

The EEG recording, pre-processing and statistical analyses were the same as in Experiment 1. ERPs time-locked to the onset of the target string presentation were obtained only for the trials followed by a correct behavioral response. RTs that were more than 2.5 SDs away from the mean of each condition and each participant were also excluded from the analysis of the response latencies. A by-subject repeated-measures ANOVA was conducted on the accuracy scores and the RTs including Group (Deaf, Hearing) as a between-subject factor, and Prime Type (Same, TL, RL) as a within-subject factor.

For the EEG data, a repeated-measures ANOVA was conducted on the average ERP amplitude similarly to Experiment 1 (same time windows, topographical factors, follow-up analyses and corrections). Apart from the topographical factors, the ERP analysis performed here included Group (Deaf, Hearing) as a between-subject factor and Prime Type (Same, TL, RL) as a within-subject factor. LME models for accuracy, RTs and ERPs led to similar results and are reported in full in the Supplementary materials. To make sure that our findings were not affected by baseline artifacts additional ANOVAs were run on the ERPs time-locked to the prime onset. These analyses led to similar results and are reported in full in the Supplementary materials.

#### Behavioral results

##### Accuracy

Deaf readers were slightly more accurate than hearing controls (Group: *F(1,38)*=6.848, *p*<.05). The Prime Type effect was significant (Prime Type: *F(2,76)*=64.624, *p*<.001), indicating that TL was less accurate than the other two conditions (TL vs Same: *t(39)*=7.735, *p*<.001; TL vs RL: *t(39)*=9.377, *p*<.001; see Figure 4). The Group x Prime Type interaction was not significant.

**Figure 3.**
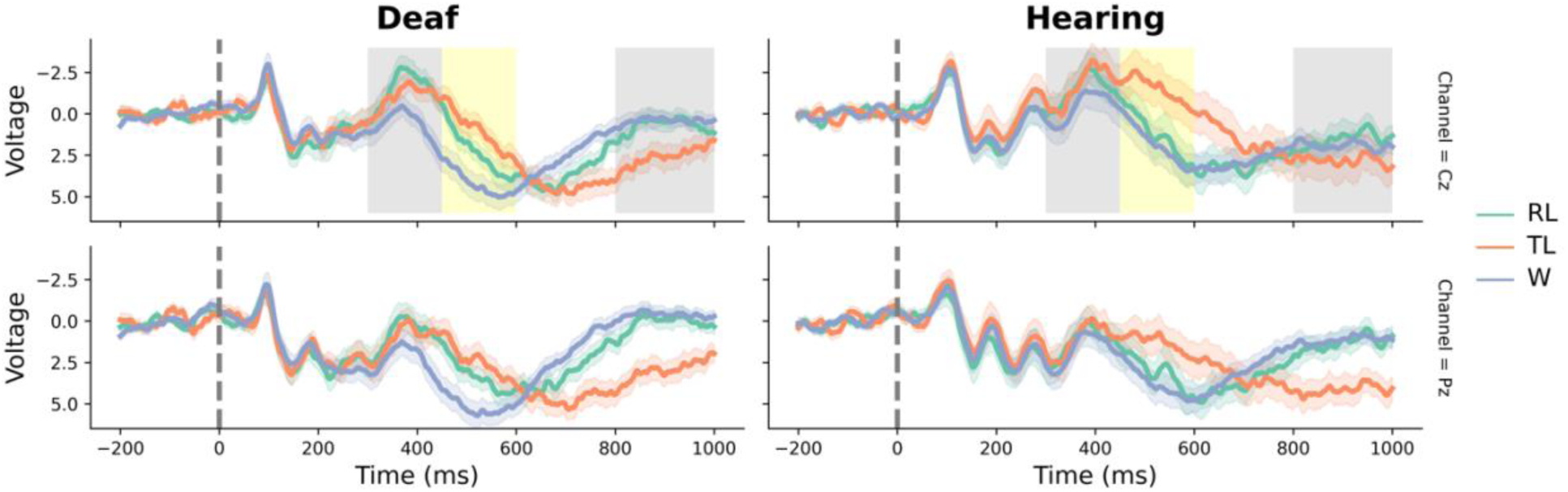
ERP waveforms to a single word study with a lexical decision task for the Deaf and Hearing groups. Shaded areas represent +/- 1SE. Cz and Pz are shown in the upper and lower panel, respectively. Each waveform represents one of the following experimental conditions: Replaced letter pseudoword (RL; e.g. tehékono), Transposed letter pseudoword (TL; e.g., tefélono), Word (W; e.g., teléfono, *phone* in Spanish). Both groups showed a greater N400 for the two pseudoword conditions (TL and RL) as compared to the Word condition. In addition, both groups showed a greater Late Positive Component (LPC) for the TL as compared to the other two conditions (RL and W). However, the RL condition showed a longer lasting N400 effect for the Deaf group as compared to hearing controls. Gray boxes highlight the time windows of the ERP effects shared across groups. Yellow boxes highlight the time window where ERP group differences were consistently reported in ANOVAs and LME models.

**Figure 4.**
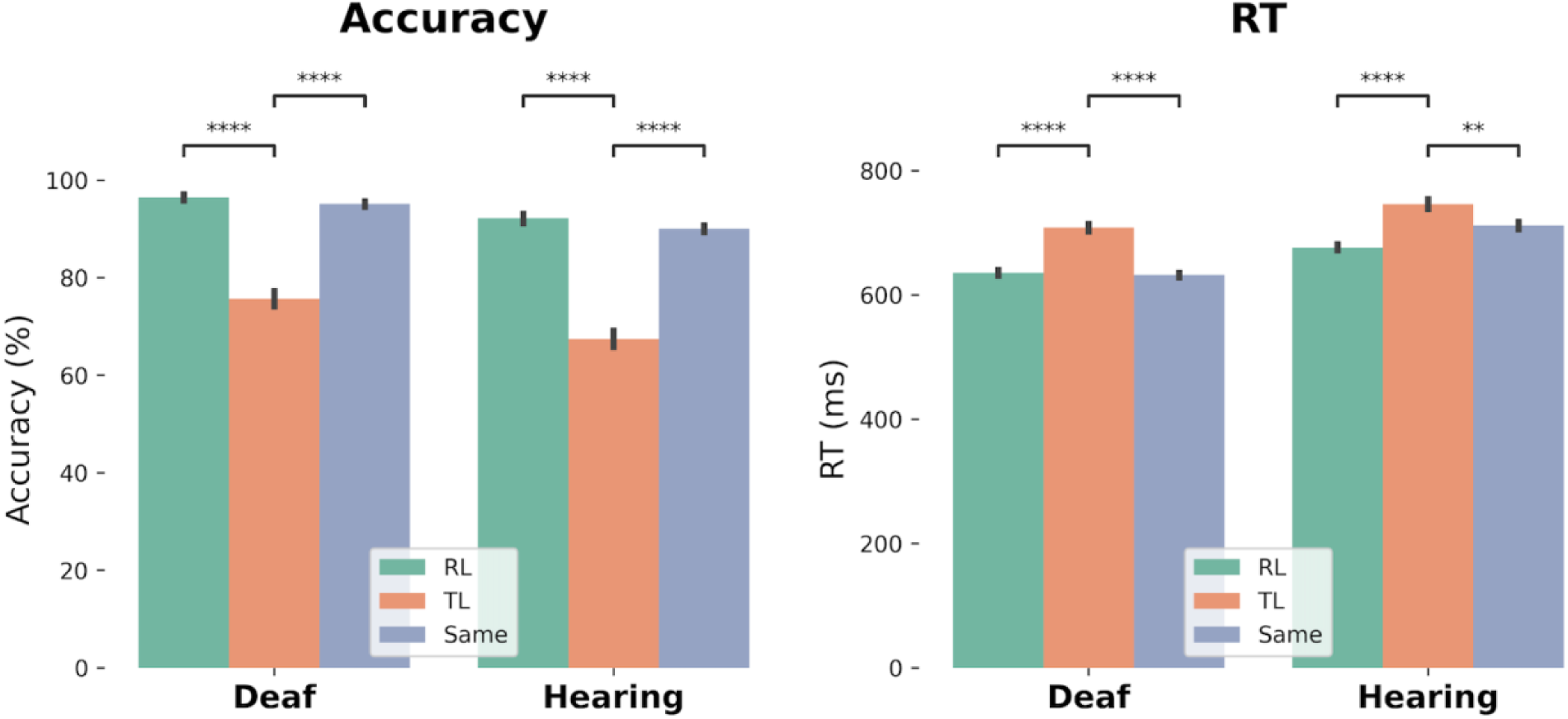
Average accuracy and reaction times for each Group and Prime Type. Error bars represent +/- 1 SE. TL was the condition showing the lowest accuracy and the slowest responses. **: p<0.01; ****: p<0.0001.

##### RT

A Prime Type effect was observed (Prime Type: *F(2,76)*=24.533, *p*<.001), suggesting that TL was the slowest condition (TL vs Same: *t(39)*=4.415, *p*<.001; TL vs RL: *t(39)*=8.128, *p*<.001; see Figure 4). No Group effect or interaction was observed.

##### ERP results

No significant effects involving our experimental conditions were found before 300 ms.

###### 300-450 ms

A N400 effect was observed for the non-identical conditions (RL and TL) as compared to the Same condition (Prime Type: *F(2,76)*=7.168, *p*<.01; Prime Type x Laterality: *F(4,152)*=3.954, *p*<.01; Prime Type x Anteriority x Laterality: *F(8,304)*=4.353, *p*<.001) and the effect reached its maximum over right posterior sensors (*ts*>3, *ps*<.01). A significant interaction Group x Prime Type x Anteriority (*F(4,152)*=2.798, *p*<.05) highlighted that this posterior effect was present only for deaf readers (RL/TL vs Same for deaf readers: *ts*>3, *ps*<.05).

###### 450-600 ms

In both groups the RL condition showed a greater posterior positivity than the other two conditions (Prime Type: *F(2,76)*=5.971, *p*<.01; Prime Type x Anteriority: *F(4,152)*=3.408, *p*<.05; RL differed from the TL and Same conditions over posterior sensors: *ts*>2, *ps*<.05).

###### 600-800 ms

In both groups, the non-identical conditions (RL and TL) showed a greater positivity as compared to the Same condition (Prime Type: *F(2,76)*=24.395, *p*<.001; Prime Type x Anteriority: *F(4,152)*=5.802, *p*<.01; Prime Type x Laterality: *F(4,152)*=2.990, *p*<.05; Prime Type x Anteriority x Laterality: *F(8,304)*=9.602, *p*<.001). This effect reached its maximum over central sensors (*ts*>5, *ps*<.001).

###### 800-1000 ms

The TL condition elicited a greater posterior positivity as compared to the other two conditions (Prime Type: *F(2,76)*=9.704, *p*<.001; Prime Type x Anteriority x Laterality: *F(8,304)*=5.624, *p*<.001; TL differed from RL and Same in central, *ts*>2, *ps*<.05, and posterior sites, *ts*>3, *ps*<.05). Critically, this posterior effect was present only in deaf readers (Group x Prime Type x Anteriority: *F(4,152)*=3.198, *p*<.05; deaf effect over central-posterior sites: *ts*>3, *ps*<.05).

#### Discussion

Experiment 2 tested the relative weight of letter position and letter identity during the processing of consonant strings in skilled deaf readers and hearing controls. Similarly to Experiment 1, behavioral results showed high levels of accuracy in both groups, with TL being the most difficult condition (i.e., the slowest and least accurate, as in Chambers, 1979; Fariña et al., 2017; O’Connor & Forster, 1981). ERPs revealed that these high levels of accuracy were achieved through different strategies. Deaf readers showed an earlier orthographic mismatch detection as compared to skilled readers, possibly due to a finer orthographic sensitivity to letter identity and letter position (in line with Gutierrez-Sigut et al., 2022; Holcomb et al., 2024). In addition, both groups differed in the later phases of re-analysis (indexed by the LPC), where letter position had a longer-lasting impact only for deaf readers.

In both Experiment 1 and 2 we have seen that, although reading accuracy is similar, the time course of orthographic analysis differs between deaf readers and controls. Specifically, a consistent group difference has been reported in the lexical/sub-lexical analysis of text (indexed by N400 effects) where deaf readers showed a consistently stronger sensitivity to letter identity. The two groups also differ in the way they process letter position, but these effects were less consistent across the two studies and mainly involved later stages of re-analysis. To deepen our understanding of the present findings, Experiment 3 and 4 will focus on deaf readers’ early sensitivity to letter identity and test whether this effect is modulated by letters’ visual features (i.e., letter visual similarity and letter case).

### Experiment 3

Experiment 3 tested the role of letter identity and visual similarity during the orthographic processing of skilled deaf readers and hearing controls. The experimental paradigm was the same as the one used in Experiment 2, where identical and non-identical prime-target pairs were presented. In Experiment 3, non-identical primes were always created through letter replacements (RL). To deepen our understanding on the role of letter visual similarity, the RL condition was further divided into letter replacements of visually similar letters and letter replacements of visually dissimilar letters.

#### Methods

##### Participants

The participants were the same as those in Experiment 1.

##### Materials

As in Experiment 2, the stimulus pairs consisted of two character strings of four letters: a prime and a target. The uppercase and lowercase version of the consonants b/B, c/C, d/D, f/F, g/G, h/H, j/J, k/K, l/L, m/M, n/N, p/P, q/Q, r/R, s/S, t/T, v/V, w/W, x/X, y/Y and z/Z were used in the strings. The prime was either identical to the target (matched item) or different to the target (mismatched item). Mismatched primes were created by modifying the internal characters of the reference string, by substituting in either visually similar characters (e.g., BWRC-BMPC, zdqp-zbgp), or dissimilar characters (e.g., BLVC-BMPC, zflp-zdgp). The similarity of the characters was controlled through the similarity scores (on a scale of 1 to 5) published by Boles and Clifford (1989). For the visually similar letters, the mean value of similarity was 3.09 (range=2.17-4.25), while for the visually dissimilar letters, the mean value was 1.93 (range=1.42-2.59). No item repetition occurred within the experiment other than that each target string appeared twice (once with a matching prime and once with a mismatching prime). Two lists were constructed with an equal number of similar and dissimilar replaced-letter primes, and such that across both lists, a given target had a mismatch with a similar-letter prime in one list and with a dissimilar-letter prime in the other list. In each condition, half the items were uppercase and half were lowercase. Participants were randomly assigned to one of the two lists. In total, each participant completed 160 trials: 80 trials with matched strings and 80 trials with mismatched strings (40 similar-letter primes and 40 dissimilar-letter primes).

##### Procedure

The procedure was the same as in Experiment 2.

##### EEG recording and analysis

The EEG recording, pre-processing and statistical analysis were the same as in Experiment 1 and 2. RTs that were more than 2.5 SDs away from the mean of each condition and each participant were also excluded from the analysis of the response latencies. A by-subject repeated-measures ANOVA was conducted on the accuracy scores and the RTs including Group (Deaf, Hearing) as a between-subject factor, and Prime Type (Same, Different but similar - Diff sl, Different and dissimilar - Diff dl) as a within-subject factor.

For the EEG data, a repeated-measures ANOVA was conducted on the average ERP amplitude similarly to Experiment 1 and 2 (same time windows, topographical factors, follow-up analyses and corrections). The ERP analysis included Group (Deaf, Hearing) as a between-subject factor and Prime Type (Same, Different but similar, Different and dissimilar) as a within-subject factor. LME models for accuracy, RTs and ERPs led to similar results and are reported in full in the Supplementary materials. Additional ANOVAs run on the ERPs time-locked to the prime onset led to similar results and are reported in full in the Supplementary materials.

#### Behavioral results

##### Accuracy

A Prime Type effect was observed (Prime Type: *F(2,76)*=12.370, *p*<.001), suggesting that mismatching primes with different letters showed the highest accuracy (Diff dl vs Diff sl: *t(39)*=5.871, *p*<.001; Diff dl vs Same: *t(39)*=4.278, *p*<.001; see Figure 6). No Group effect or interaction was observed.

**Figure 5.**
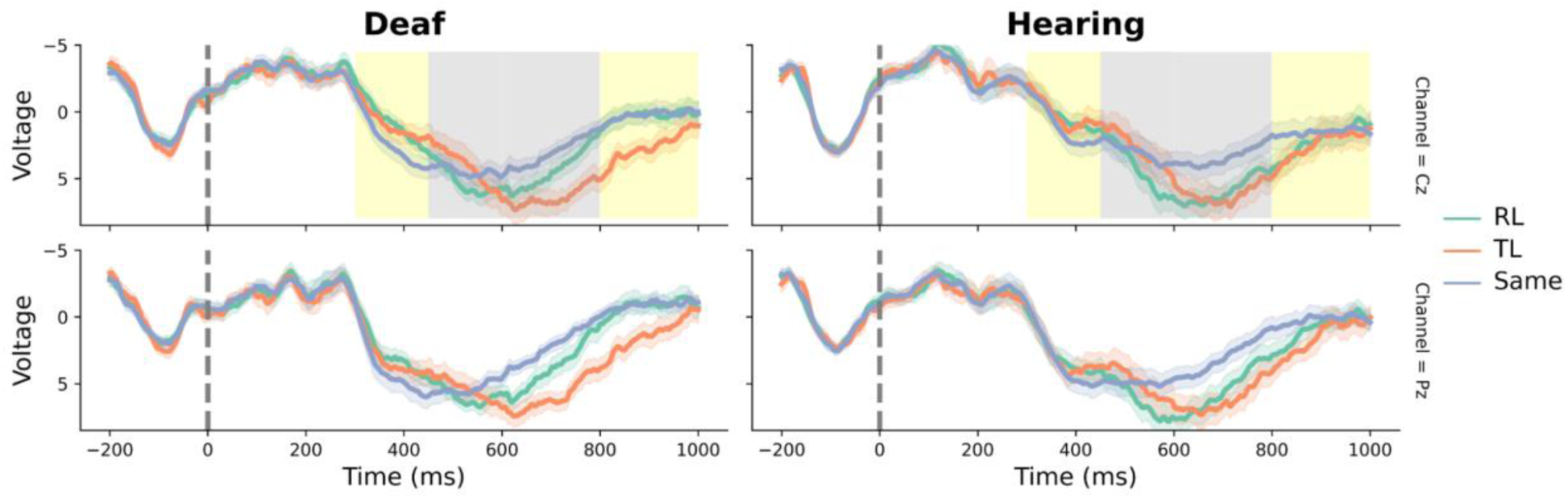
ERP waveforms of a priming study with a same-different decision task manipulating letter position and identity for the Deaf and Hearing groups. Shaded areas represent +/- 1SE. Cz and Pz are shown in the upper and lower panel, respectively. Each waveform represents one of the following experimental conditions: Replacement of letters (RL; e.g. NFLT-NDXT), Transposition of letters (TL; e.g., NXDT-NDXT), Same (e.g., NDXT-NDXT). Both groups showed a greater LPC for the two non-identical conditions (TL and RL) as compared to the Same condition. Gray boxes highlight the time windows of the ERP effects shared across groups. Two group differences were observed (highlighted by the yellow boxes). There was a greater N400 for non-identical conditions (TL and RL) as compared to the Same condition only for the skilled deaf readers. In addition, in the late time window, deaf readers showed a long-lasting LPC effect for the TL condition.

**Figure 6.**
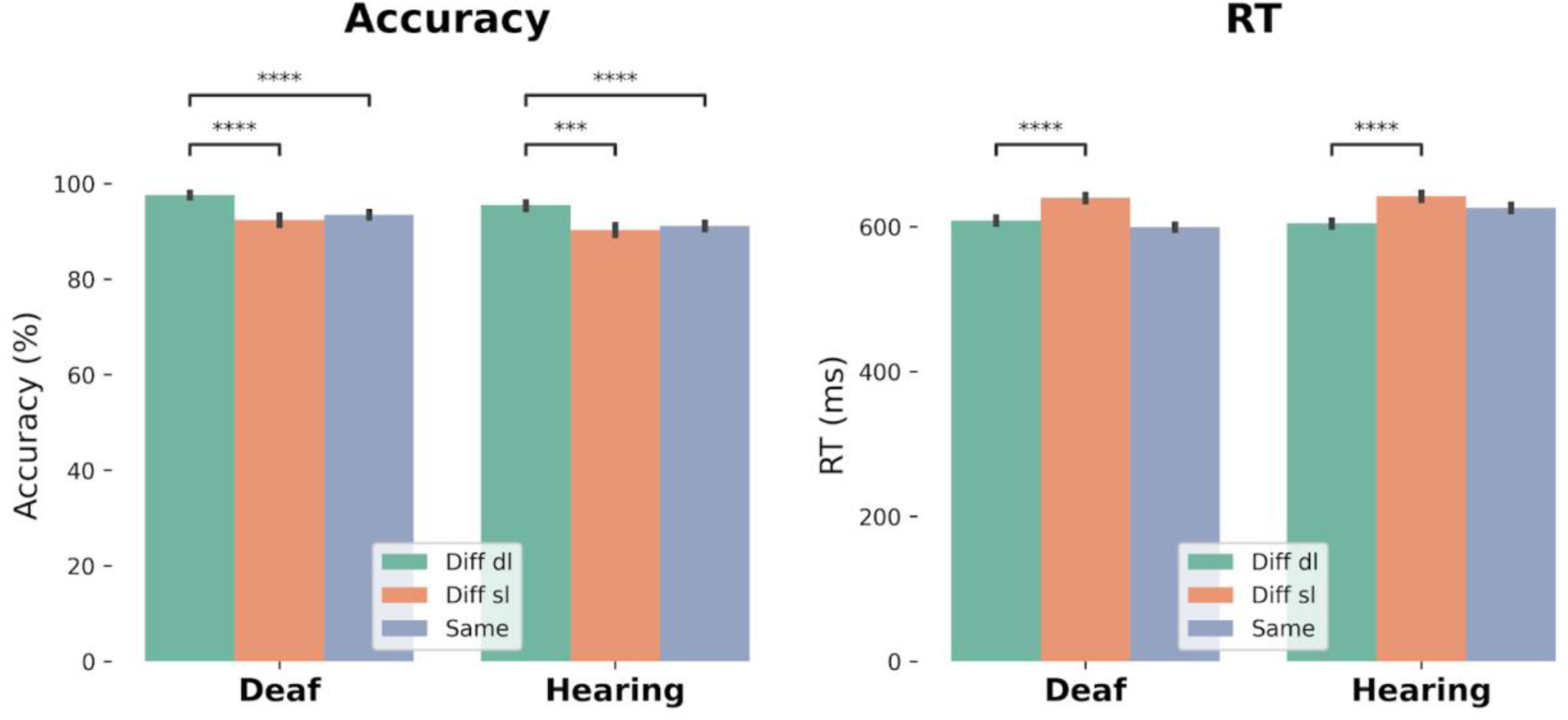
Average accuracy and reaction times for each Group and Prime Type. Error bars represent +/- 1 SE. Both groups were slower and less accurate at detecting differences when the strings showed visually similar as compared to different letters. ***: p<0.001; ****: p<0.0001.

**Figure 7.**
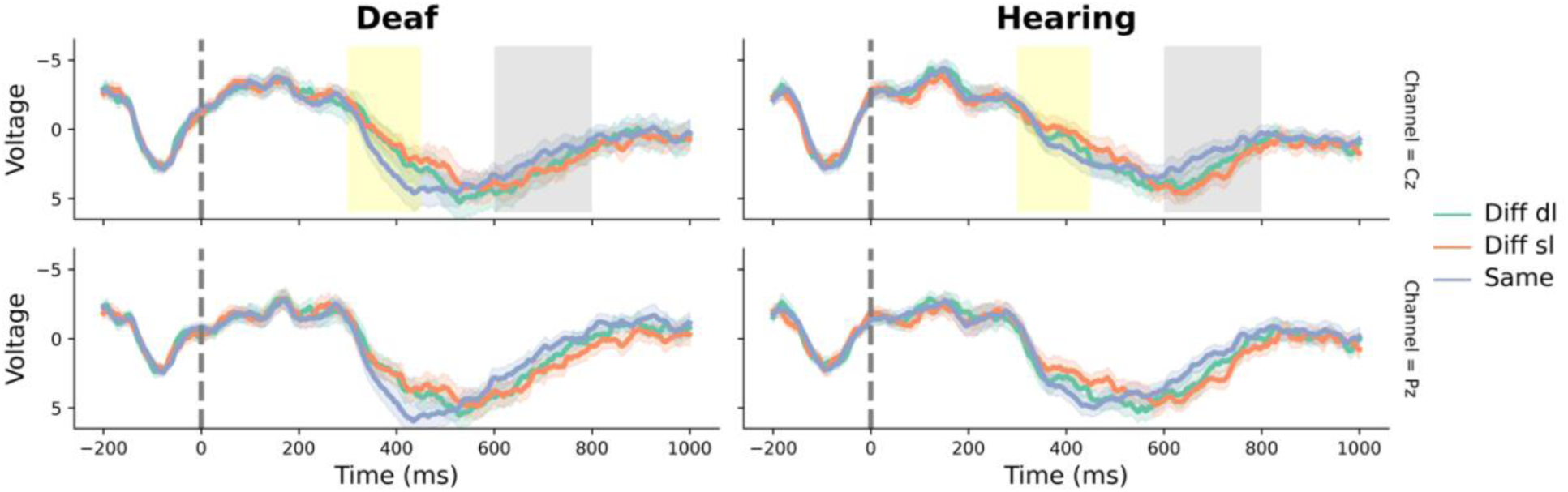
ERP waveforms of a priming study with a same-different decision task manipulating visual similarity of letters for the Deaf and Hearing groups. Shaded areas represent +/- 1SE. Cz and Pz are shown in the upper and lower panel, respectively. Each waveform represents one of the following experimental conditions: Different strings with different letters (Diff dl; e.g. LNKD-LRSD), Different strings with visually similar letters (Diff sl; e.g., LPZD-LRSD), Same strings (e.g., LRSD-LRSD). Gray boxes highlight the time window of the ERP effects shared across groups: a greater LPC for the two non-identical conditions (Diff dl and Diff sl) as compared to the Same condition. Yellow boxes highlight the time window where ERP group differences were reported: only the Deaf group showed a greater N400 effect for the two non-identical conditions (Diff dl and Diff sl) as compared to the Same condition.

##### RT

A Prime Type effect was observed (Prime Type: *F(2,76)*=7.808, *p*<.01), suggesting that mismatching primes with different letters showed faster responses as compared to those with similar letters (Diff sl vs Diff dl: *t(39)*=9.147, *p*<.001; see Figure 6). No Group effect or interaction was observed.

##### ERP results

No significant effects involving our experimental conditions were found before 300 ms.

###### 300-450 ms

A N400 effect was observed for the non-identical conditions (Diff dl and Diff sl) as compared to the same condition (Prime Type: *F(2,76)*=7.248, *p*<.01), which was right-lateralized (Prime Type x Laterality: *F(4,152)*=6.103, *p*<.001; Diff dl/Diff sl vs same: *ts*>3.5, *ps*<.01). A marginal interaction Group x Prime Type x Laterality (*F(4,152)*=2.090, *p*=.052) highlighted that this right-lateralized effect was present only in the deaf readers (deaf: *ts*>3, *ps*<.05; controls: *ts*<2.4).

###### 450-600 ms

None of the contrasts of interest was significant.

###### 600-800 ms

In both groups, the non-identical conditions (Diff dl and Diff sl) showed a greater positivity as compared to the same condition (Prime Type: *F(2,76)*=15.243, *p*<.001). This effect reached its maximum over left and right central sensors (*ts*>4, *ps*<.01).

###### 800-1000 ms

None of the contrasts of interest was significant.

#### Discussion

Experiment 3 examined the role of letter identity and letter visual similarity during the orthographic analysis of consonant strings in deaf readers and controls. Similarly to Experiment 2, a priming design was employed where prime and target could be identical or not (RL). To further improve our understanding on the alternative reading strategies used by skilled deaf readers, letter similarity was also manipulated so that the replaced letters could be visually similar or dissimilar to the original target letters. In line with our previous findings, deaf readers and hearing controls differed in the way they processed letter replacements, with deaf readers showing an earlier detection of prime-target orthographic matches. This group difference was present regardless of the degree of visual similarity of the mismatching letters, suggesting that the refined sensitivity to letter identity of skilled deaf readers does not seem to depend on visual letter form. Experiment 4 further tested this hypothesis by manipulating an additional dimension of letters’ visual appearance: letter case.

### Experiment 4

Experiment 4 investigates the impact of letter case on the time course of orthographic processing in deaf readers and matched controls. The experimental paradigm employed was the same as the one used in Experiment 2 and 3. Letter identity was again manipulated so that prime and target could be identical or not (RL). Letter case was also manipulated so that the prime had the same or different case compared to the target.

#### Methods

##### Participants

The participants were the same as those in Experiment 1.

##### Materials

As in Experiment 2 and 3, the stimulus pairs consisted of two character strings of four letters: a prime and a target. The uppercase and lowercase version of the consonants b/B, c/C, d/D, f/F, g/G, h/H, j/J, k/K, l/L, m/M, n/N, p/P, q/Q, r/R, s/S, t/T, v/V, w/W, x/X, y/Y and z/Z were used in the strings. The prime either had the same letters as the target (matched item) or different letters target (mismatched item). The first and last letters of the strings were always uppercase; the case of the middle two letters was manipulated as follows. The matched primes could be identical in case to the target (e.g., STFV-STFV, MdpJ-MdpJ) or in a different case (e.g., StfV-STFV, MDPJ-MdpJ). Mismatched primes had different internal characters either in the same case as the target (e.g., SKLV-STFV, MthJ-MdpJ) or in a different case (e.g., SklV-STFV, MTHJ-MdpJ). The visual similarity between replaced letters was kept low using the similarity values (on a scale of 1 to 5) published by Boles and Clifford (1989). For the replaced letter condition, the mean value of similarity was 2.67 (range=1.54-4.05), while for the same letter condition the mean was 4.14 (range=2.16-5.00). No item repetition occurred within the experiment. Two lists were constructed with an equal number of matched and mismatched items, half of which had the same case, the other half had different cases; across both lists, a given target had a matching prime in one list and a mismatching prime in the other list. In each condition, half the target strings had uppercase letters and half had internal letters in lowercase. Participants were randomly assigned to one of the two lists. In total, each participant completed 160 trials: 80 trials with matched strings (40 same case and 40 different case) and 80 trials with mismatched strings (40 same case and 40 different case).

##### Procedure

The procedure was the same as in Experiment 2 and 3. Importantly, participants were instructed to decide if the character strings contained the same letters, regardless of case.

##### EEG recording and analysis

The EEG recording, pre-processing and statistical analysis were the same as in Experiment 1, 2, and 3. RTs that were more than 2.5 SDs away from the mean of each condition and each participant were also excluded from the analysis of the response latencies. A by-subject repeated-measures ANOVA was conducted on the accuracy scores and the RTs including Group (Deaf, Hearing) as a between-subject factor, and Prime Type (Matching-same case, Matching-different case, Mismatch-same case, Mismatch-different case) as a within-subject factor.

For the EEG data, a repeated-measures ANOVA was conducted on the average ERP amplitude within the following time windows: 300-450 ms, 450-600 ms, 600-800 ms and 800-1000 ms. This analysis included Group (Deaf, Hearing) as a between-subject factor and Prime Type (Same sc: Matching-same case, Same dc: Matching-different case, Diff sc: Mismatching-same case, Diff dc: Mismatching-different case) as a within-subject factor. LME models for accuracy, RTs and ERPs led to similar results and are reported in full in the Supplementary materials. Additional ANOVAs run on the ERPs time-locked to the prime onset led to similar results and are reported in full in the Supplementary materials.

#### Behavioral results

##### Accuracy

No effect reached significance.

##### RT

There was a Prime Type effect (Prime Type: *F(2,76)*=7.343, *p*<.001), suggesting that when the strings were exactly the same (i.e., Same strings with the same case, Same sc) RTs were faster compared to all other conditions (Same sc vs. Same dc: *t(39)*=6.694, *p*<.001; Same sc vs. Diff sc: *t(39)*=2.964, *p*<.001; Same sc vs. Diff dc: *t(39)*=3.176, *p*<.001). The Group x Prime Type was also significant (Group x Prime Type: *F(3,114)*=2.870, *p*<.05), indicating that these RTs differences were more evident in deaf readers (deaf readers: Same sc vs. Same dc: *t(39)*=7.762, *p*<.001; Same sc vs. Diff sc: *t(39)*=4.288, *p*<.001; Same sc vs. Diff dc: *t(39)*=5.110, *p*<.001; hearing controls: Same sc vs. Same dc: *t(39)*=3.391, *p*<.001; see Figure 8).

**Figure 8.**
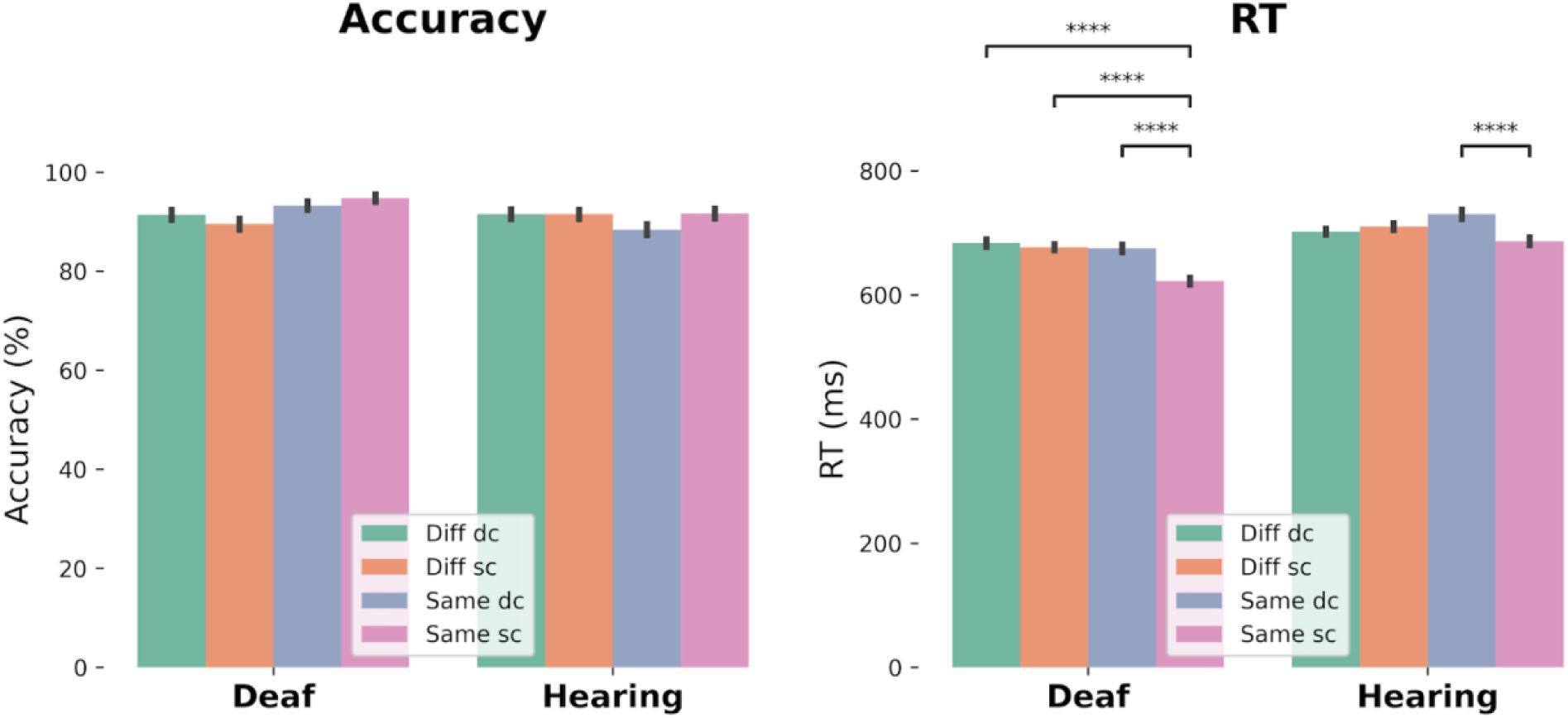
Average accuracy and reaction times for each Group, Prime Type and Case. Error bars represent +/- 1 SE. The identity condition (i.e., where the strings were the same and had the same case) was the fastest and this was more evident in the deaf reader group. ****: p<0.0001.

##### ERP results

No significant effects involving our experimental conditions were found before 300 ms.

###### 300-450 ms

Both groups showed an N400 effect for the non-identical conditions (Diff dc and Diff sc) as compared to the same conditions (Same dc and Same sc; Prime Type: *F(1,38)*=13.269, *p*<.001), which reached its maximum over right posterior sensors (Prime Type x Anteriority: *F(2,76)*=6.002, *p*<.05; Prime Type x Laterality: *F(2,76)*=4.593, *p*<.05; right sensors, *ts*>3.5, *ps*<.01, posterior sensors, *ts*>4, *ps*<.01). This effect was stronger and more widely distributed for deaf readers (Group x Prime Type: *F(1,38)*=4.363, *p*<.05; Group x Prime Type x Laterality: *F(2,76)*=3.500, *p*<.05; the deaf group showed a difference over medial and right sites: *ts*>3, *ps*<.01, while the hearing controls showed a difference only over right sites: *ts*>2, *ps*<.01).

###### 450-600 ms

An N400 effect persisted over central-posterior sensors (Prime Type: *F(1,38)*=14.298, *p*<.001; Prime Type x Anteriority: *F(2,76)*=13.505, *p*<.001, central-posterior sensors: *ts*>3, *ps*<.01). This effect was still more widely distributed in deaf readers as compared to hearing controls (Group x Prime Type x Anteriority: *F(2,76)*=8.548, *p*<.01; deaf showed an effect over central and posterior sensors, *ts*>2, *ps*<.05, while in hearing controls the effect was only over posterior sensors, *ts*>2, *ps*<.01).

###### 600-800 ms

Both groups showed a greater positivity for the non-identical conditions (Diff dc and Diff sc) as compared to the same conditions (Prime Type x Anteriority x Laterality: *F(4,152)*=3.157, *p*<.05). This effect was more widely distributed for the same case strings (Prime Type x Case: *F(1,38)*=9.691, *p*<.01; Prime Type x Case x Anteriority: *F(2,76)*=21.590, *p*<.001; same cases showed a difference over central posterior sensors, *ts*>2, *ps*<.05, while different cases showed a difference only over posterior sensors, *ts*>2, *ps*<.05).

###### 800-1000 ms

The central posterior positive effect persisted only for the same case string (Prime Type x Case x Anteriority: *F(2,76)*=11.978, *p*<.001; central and posterior sensors, *ts*>2, *ps*<.05).

#### Discussion

Experiment 4 tested the role of letter identity and letter case in the orthographic processing of skilled deaf readers and hearing controls. Behavioral results again showed high levels of accuracy for both groups, with the identical condition being the fastest. As compared to hearing controls, deaf readers were faster at spotting pure visual identity among all other conditions. In the ERPs, both groups showed an N400 and an LPC effect for non-identical as compared to the identical conditions in line with previous findings (Emmorey et al., 2021; Mitra & Coch, 2009). Consistent with our previous Experiments, deaf and hearing readers differed in the way they processed the identity of orthographic information. Skilled deaf readers showed a stronger and more distributed N400 effect for letter replacements as compared to hearing controls. This group difference was present regardless of visual similarity between prime and target, indicating that skilled deaf readers have a refined early sensitivity to abstract, orthographic letter identity which does not depend on the letters’ visual form.

## General Discussion

Four EEG and behavioral experiments examined how deaf readers and hearing controls process orthographic information within lexical (Exp. 1) and sub-lexical (Exp. 2-4) domains. Across the four experiments both groups showed similarly accurate performance. However, the EEG response revealed a consistent difference between deaf and hearing readers in the lexical and sub-lexical analysis of letter identity around 400 ms and in the later re-analysis phase of letter position (Table 2).

**Table 2.**
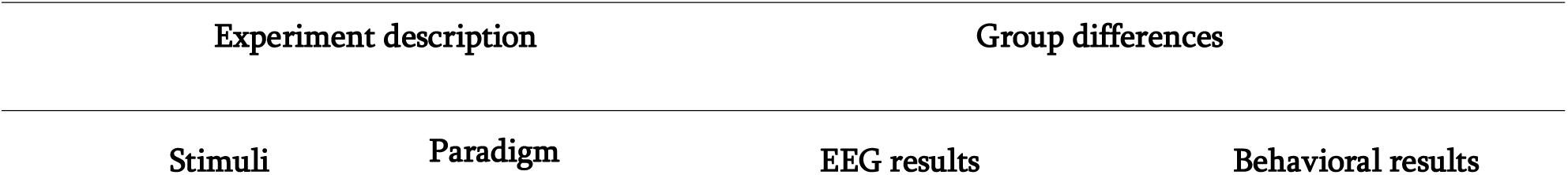
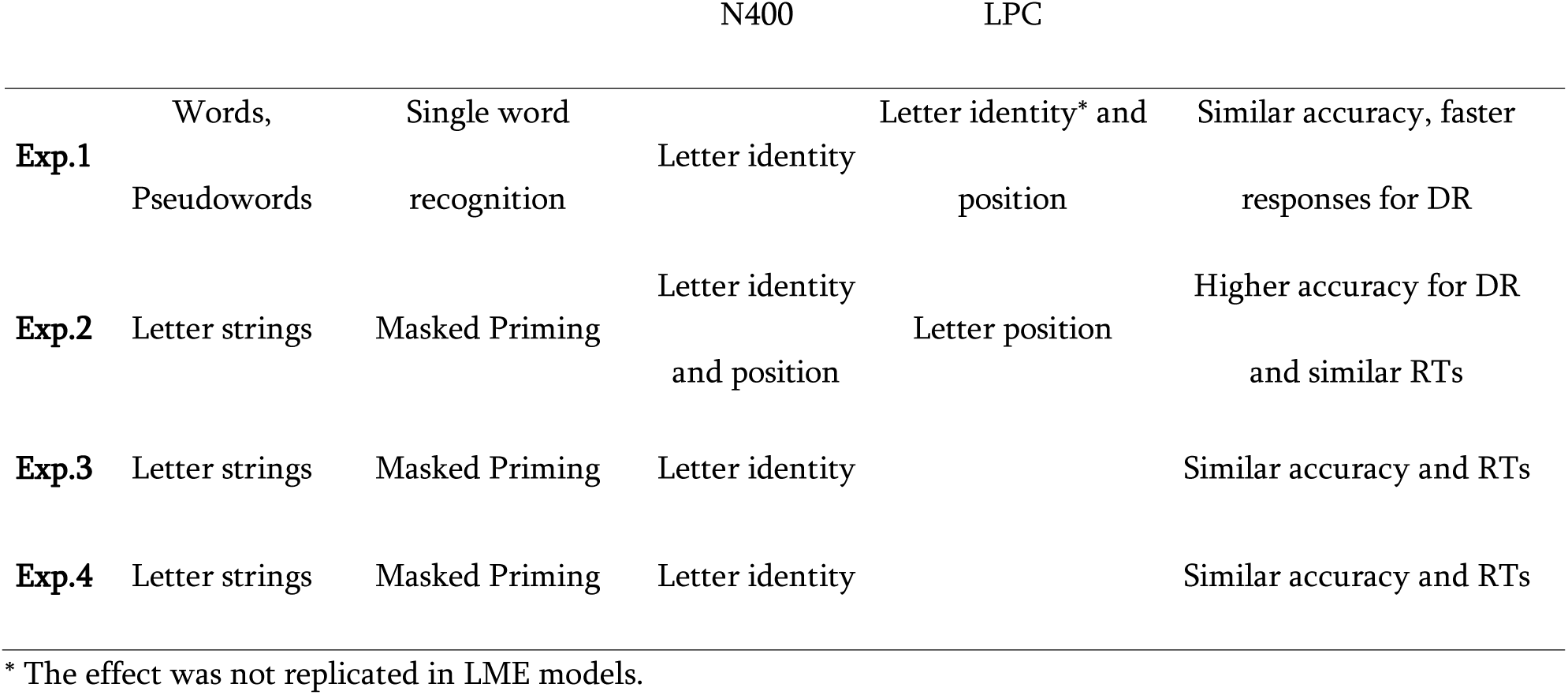
Overview of the group differences observed in EEG and behavioral results for each experiment. EEG results list the orthographic factors (i.e. letter identity, letter position) showing significant group differences. DR: Deaf readers

Specifically, within a lexical domain (Exp. 1), deaf readers show longer lasting N400 effects for changes in letter identity as compared to hearing controls. In the Experiments investigating the sub-lexical domain through letter strings (Exp. 2-4), deaf readers show earlier and/or greater N400 effects for changes in letter identity as compared to hearing controls, which were observed regardless of variations to letter visual form (i.e. visual similarity and case; Exp. 3-4). Where letter position was manipulated (Exp.1-2), a group difference was consistently observed in the later phase of re-analysis with earlier/ longer-lasting LPC effects (Exp.1-2) for letter transpositions in deaf readers.

This suggests that deaf readers and hearing controls can reach similar high levels of accuracy in text recognition by leveraging different strategies of orthographic processing. As compared to hearing controls, deaf readers consistently show a higher sensitivity to letter identity around 400 ms after stimulus presentation (Figure 10) and a more pronounced sensitivity to letter position in a later phase of orthographic re-analysis.

**Figure 9.**
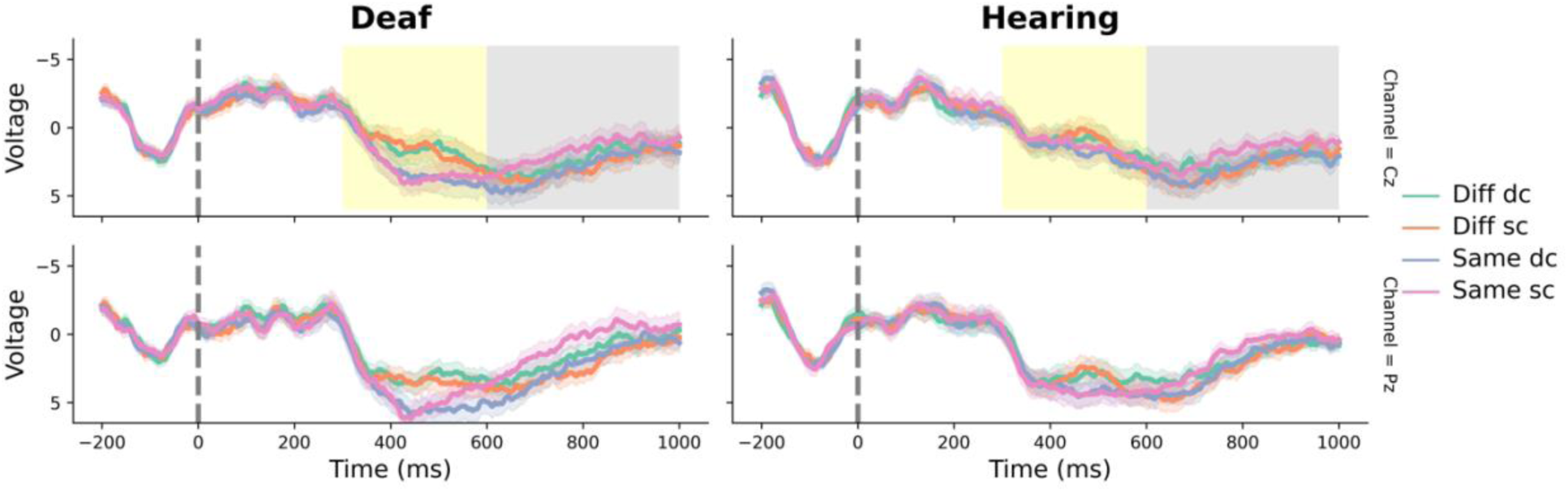
ERP waveforms of a priming study with a same-different decision task manipulating letter case for the Deaf and Hearing groups. Shaded areas represent +/- 1SE. Cz and Pz are shown in the upper and lower panel, respectively. Each waveform represents one of the following experimental conditions: Different strings with different cases (Diff dc; e.g. SmfV - SGTV), Different strings with same case (Diff sc; e.g., SMFV - SGTV), Same strings with different cases (Same dc; e.g. SgtV - SGTV), Same strings with same case (Same sc; e.g., SGTV - SGTV). Both groups showed a greater N400 and LPC for the different strings as compared to the same strings with matching cases. Gray boxes highlight the time window of the ERP effects shared across groups. Yellow boxes highlight the time window where ERP group differences were reported: the N400 effect for different strings relative to the same strings (regardless of case) was stronger for the deaf compared to the hearing group.

**Figure 10.**
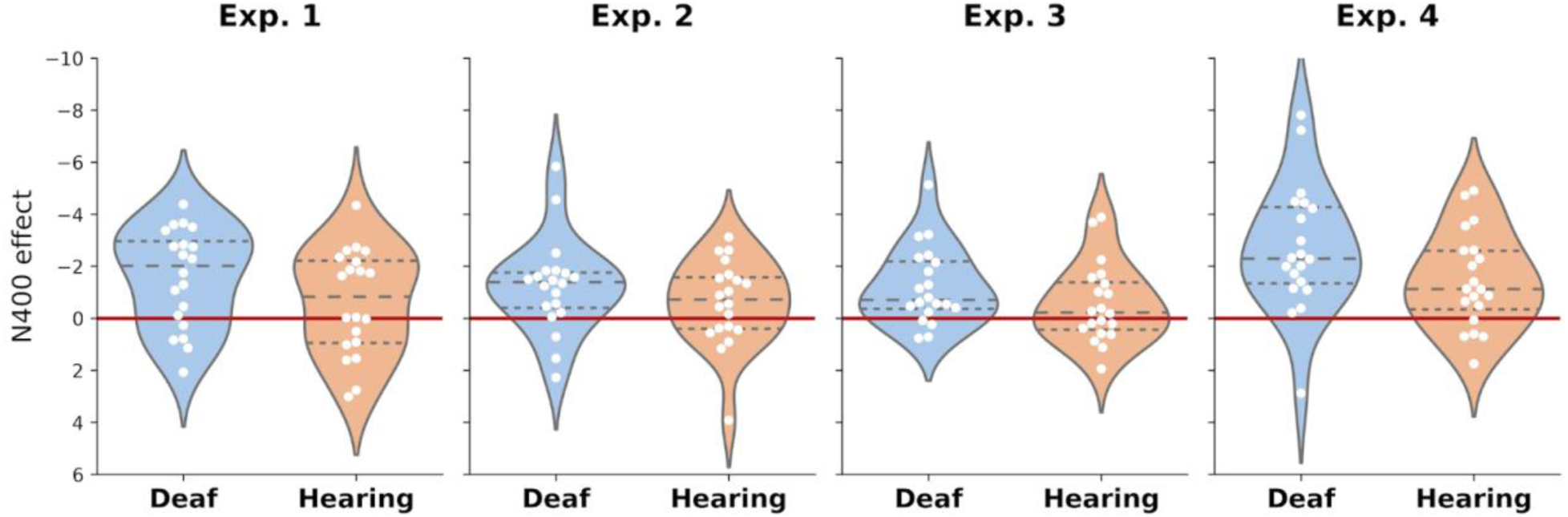
Overview of the individual N400 effects for letter replacement. Data points are displayed for each group and experiment in the time window where significant group differences were observed. Deaf readers’ distribution is consistently shifted towards stronger N400 effects as compared to hearing controls’ distribution.

These findings add to the present literature by showing that proficient deaf readers not only have a different phonological processing of text as compared to hearing controls (Costello et al., 2021), but they also show a distinct time course of orthographic processing (Bélanger & Rayner, 2015; Emmorey et al., 2016; Gutierrez-Sigut et al., 2019; Lee et al., 2022; Peleg et al., 2020; cf. Fariña et al., 2017; Meade et al., 2019, 2020). Crucially, these differences do not hinder deaf readers’ text recognition accuracy (which is either similar or even slightly higher than controls, see Exp. 2) and, rather, they represent valid alternative strategies for high reading achievement. Our findings further qualify these alternative reading strategies by showing that, in early phases of text processing (which are the least affected by motor response preparation), a greater sensitivity to letter identity supports deaf readers’ proficient reading. Findings from Exp. 3 and 4 suggest that deaf readers show a refined sensitivity to visual matches which does not uniquely depend on the visual appearance of graphemes (i.e., letter visual similarity or case), but it is probably related to a more abstract representation of orthographic information (Bélanger & Rayner, 2015) and/or a greater reliance on orthographic cues (Emmorey et al., 2016; Gutierrez-Sigut et al., 2019; Holcomb et al., 2024; Lee et al., 2022; Peleg et al., 2020). Moreover, our findings are not in line with models that assume a similar degree of precision of orthographic representations between deaf and hearing readers (Meade et al., 2019, 2020).

This experimental evidence confirms the plasticity of the reading system and the existence of alternative pathways to reading success. An absence of acoustic experience can trigger the development of an increased sensitivity to orthographic information (i.e., letter identity and letter position) during different stages of text processing. This suggests that phonological coding is not essential to have strong orthographic representations for skilled readers, which has potential educational and clinical implications.

The majority of deaf children show difficulties learning to read and lag behind their peers despite early intervention and technological aids (Allen, 1986; Chamberlain & Mayberry, 2008; Conrad, 1979; Geers, 2003; Mayberry, 2002; Miller, 2006; Musselman, 2000; Strong & Prinz, 1997; Wauters et al., 2006). Building up substantial orthographic knowledge might represent an important educational strategy to overcome delays in reading acquisition. In addition, reading disorders characterized by reduced phonological awareness even in hearing individuals, such as dyslexia, might be compensated by a reinforced orthographic knowledge (Leinonen et al., 2001).

## Limitations

Our late ERP responses should be interpreted with caution as they overlap with the time window of button press responses (and related motor preparation) and different types of responses have been contrasted. However, our most consistent finding across studies is relative to the early phase of orthographic analysis, which has reduced overlapping risks as it happens at least 200 ms earlier than button press. Hence, the most robust evidence provided by our studies supports the presence of different orthographic strategies for deaf and hearing readers, which mainly regards the analysis of letter identity, rather than letter position. Finally, it should be noted that our priming studies did not report significant effects before the N400 time window in contrast with previous research showing N250 modulations (Duñabeitia et al., 2009, 2012; Emmorey & Lee, 2021; Holcomb et al., 2024; Lee et al., 2022; Meade et al., 2020). This difference might be due to a paradigm difference as our target items were consonant strings rather than real words. This asymmetry might also be due to the degree of replicability of early ERP effects as compared to later ones, with stronger cross-paradigm consistency for N400 effects as compared to earlier potentials (Kim et al., 2024; Segalowitz & Barnes, 1993).

## Conclusions

In conclusion, our findings demonstrate that proficient deaf readers achieve high levels of reading accuracy through distinct orthographic processing strategies. Specifically, they exhibit heightened sensitivity to letter identity during early lexical and sub-lexical processing. These differences in the temporal dynamics of orthographic processing, which are independent of visual letter form, do not impede reading performance but rather reflect alternative pathways to reading success. This evidence underscores the plasticity of the reading system and highlights the potential for educational approaches that strengthen orthographic knowledge to support reading acquisition in deaf children and individuals with phonologically-based reading difficulties. Overall, these results expand our understanding of the cognitive mechanisms underlying proficient reading and provide a framework for interventions that leverage orthographic expertise as a compensatory strategy.

## Constraints on Generality paragraph

Highly proficient deaf readers provide a particularly informative population for examining alternative orthographic processing strategies when phonological processing is limited.

## Acknowledgements

The authors are grateful for assistance from CNSE (the Spanish National Body of Deaf People), various Deaf People’s Associations throughout Spain and two universities (Pompeu Fabra University in Barcelona and University of La Laguna in Tenerife), which provided facilities to run the experiments. The authors thank the study participants, without whom this work would not have been possible.

## Supplementary Materials

### S1. Mixed linear models for behavioral data

#### Exp 1

LMERs with Word Type and Group as fixed factors (reference being TL condition and deaf group, respectively) and a by-subject random intercept.

| Accuracy | $\beta$ | SE | $t$ | $p$ | CI<br>[0.025,0.975] |
| --- | --- | --- | --- | --- | --- |
| Intercept | 0.800 | 0.015 | 53.511 | <0.0001 | [0.770, 0.829] |
| Group (controls) | -0.021 | 0.016 | -1.280 | 0.20 | [-0.052, 0.011] |
| Word Type (RL) | 0.183 | 0.017 | 10.928 | <0.0001 | [0.150, 0.216] |
| Word Type (W) | 0.190 | 0.017 | 11.320 | <0.0001 | [0.157, 0.223] |
| Subject | 0.001 | 0.010 |  |  |  |

| RT | $\beta$ | SE | $t$ | $p$ | CI<br>[0.025,0.975] |
| --- | --- | --- | --- | --- | --- |
| Intercept | 766.172 | 25.909 | 25.571 | <0.0001 | [715.4, 817.0] |
| Group (controls) | 121.218 | 35.147 | 3.449 | 0.001 | [52.3,190.1] |
| Word Type (RL) | -114.802 | 12.687 | -9.049 | <0.0001 | [-139.7, -89.9] |
| Word Type (W) | -155.925 | 12.687 | -12.290 | <0.0001 | [-180.8, -131.1] |
| Subject | 11279.909 | 60.910 |  |  |  |

#### Exp 2

LMERs with Prime Type and Group as fixed factors (reference being TL condition and deaf group, respectively) and a by-subject random intercept.

| Accuracy | $\beta$ | SE | $t$ | $p$ | CI<br>[0.025,0.975] |
| --- | --- | --- | --- | --- | --- |
| Intercept | 0.744 | 0.021 | 36.074 | <0.0001 | [0.704, 0.785] |
| Group (controls) | -0.059 | 0.023 | -2.617 | 0.009 | [-0.103, -0.015] |
| Prime Type (RL) | 0.228 | 0.022 | 10.286 | <0.0001 | [0.185, 0.272] |
| Prime Type (Same) | 0.211 | 0.022 | 9.506 | <0.0001 | [0.167, 0.254] |
| Subject | 0.002 | 0.014 |  |  |  |

| RT | $\beta$ | SE | $t$ | $p$ | CI<br>[0.025,0.975] |
| --- | --- | --- | --- | --- | --- |
| Intercept | 707.050 | 26.539 | 26.642 | <0.0001 | [655.1, 759.1] |
| Group (controls) | 54.781 | 36.223 | 1.512 | 0.130 | [-16.2, 125.8] |
| Prime Type (RL) | -79.061 | 12.030 | -6.572 | <0.0001 | [-102.6, -55.5] |
| Prime Type (Same) | -60.777 | 12.030 | -5.052 | <0.0001 | [-84.4, -37.2] |
| Subject | 12156.41 | 68.232 |  |  |  |

#### Exp 3

LMERs with Prime Type and Group as fixed factors (reference being Diff dl condition and deaf group, respectively) and a by-subject random intercept.

| Accuracy | $\beta$ | SE | $t$ | $p$ | CI<br>[0.025,0.975] |
| --- | --- | --- | --- | --- | --- |
| Intercept | 0.977 | 0.010 | 94.823 | <0.0001 | [0.956, 0.997] |
| Group (controls) | -0.022 | 0.011 | -1.917 | 0.055 | [-0.044, 0.0001] |
| Prime Type (Diff sl) | -0.052 | 0.011 | -4.734 | <0.0001 | [-0.074, -0.031] |
| Prime Type (Same) | -0.043 | 0.011 | -3.861 | <0.0001 | [-0.065, -0.021] |
| Subject | 0.0001 | 0.007 |  |  |  |

| RT | $\beta$ | SE | $t$ | $p$ | CI<br>[0.025,0.975] |
| --- | --- | --- | --- | --- | --- |
| Intercept | 601.237 | 21.036 | 28.581 | <0.0001 | [560.1, 642.5] |
| Group (controls) | 10.926 | 28.866 | 0.379 | 0.705 | [-45.7, 67.5] |
| Prime Type (Diff sl) | 33.263 | 8.816 | 3.773 | <0.0001 | [16.0, 50.5] |
| Prime Type (Same) | 9.289 | 8.816 | 1.054 | 0.292 | [-8.0, 26.6] |
| Subject | 7814.09 | 59.127 |  |  |  |

#### Exp 4

LMERs with Prime Type and Group as fixed factors (reference being Same sc condition and deaf group, respectively) and a by-subject random intercept.

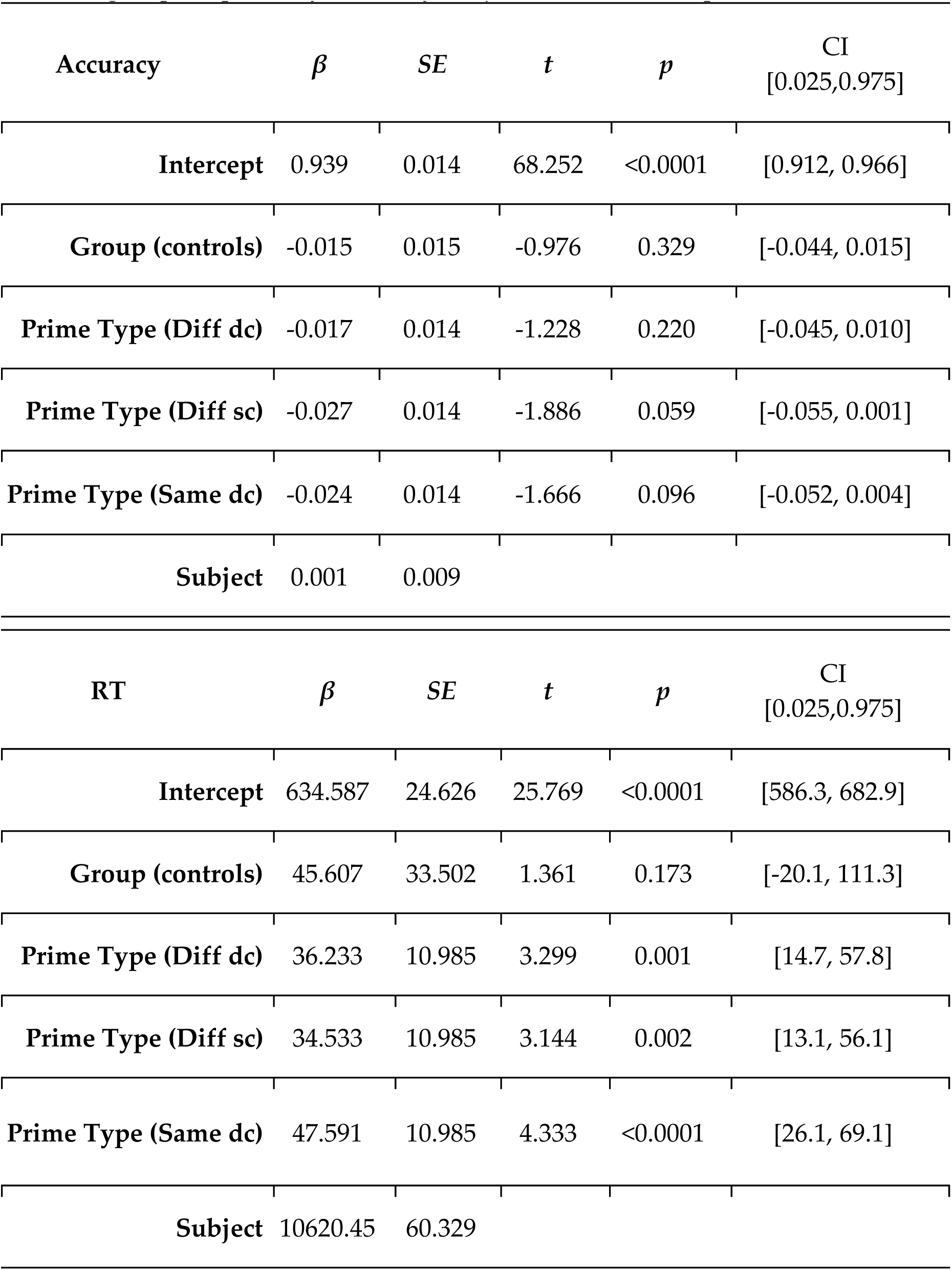

Similar LMER to the previous one including the interaction between Prime Type and Group (reference being Same sc condition and the factor Group has been releveled for follow up analyses).

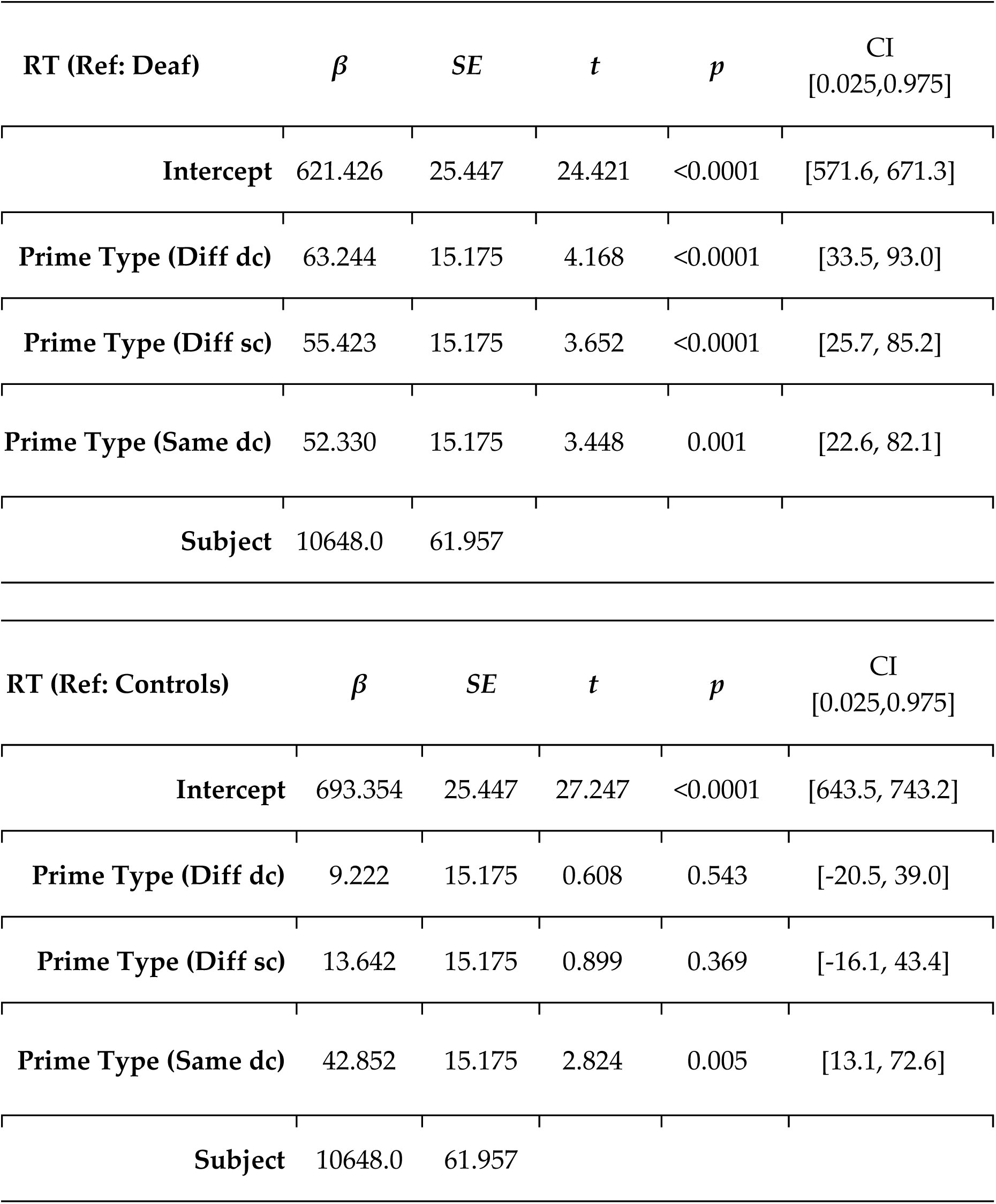

| RT (Ref: Deaf) | $\beta$ | SE | $t$ | $p$ | CI<br>[0.025,0.975] |
| --- | --- | --- | --- | --- | --- |
| **Intercept** | 621.426 | 25.447 | 24.421 | <0.0001 | [571.6, 671.3] |
| **Prime Type (Diff dc)** | 63.244 | 15.175 | 4.168 | <0.0001 | [33.5, 93.0] |
| **Prime Type (Diff sc)** | 55.423 | 15.175 | 3.652 | <0.0001 | [25.7, 85.2] |
| **Prime Type (Same dc)** | 52.330 | 15.175 | 3.448 | 0.001 | [22.6, 82.1] |
| **Subject** | 10648.0 | 61.957 |  |  |  |

| RT (Ref: Controls) | $\beta$ | SE | $t$ | $p$ | CI<br>[0.025,0.975] |
| --- | --- | --- | --- | --- | --- |
| **Intercept** | 693.354 | 25.447 | 27.247 | <0.0001 | [643.5, 743.2] |
| **Prime Type (Diff dc)** | 9.222 | 15.175 | 0.608 | 0.543 | [-20.5, 39.0] |
| **Prime Type (Diff sc)** | 13.642 | 15.175 | 0.899 | 0.369 | [-16.1, 43.4] |
| **Prime Type (Same dc)** | 42.852 | 15.175 | 2.824 | 0.005 | [13.1, 72.6] |
| **Subject** | 10648.0 | 61.957 |  |  |  |

### S2. Mixed linear models for ERPs

LMERs were performed on the average voltage from the central-posterior channels (i.e., C3, Cz, C4, P3, Pz, P4), as these were the sites showing the maximum ERP responses (i.e., N400, LPC).

#### Exp 1

LMERs with Word Type and Group as fixed factors (reference being W condition) and a by-subject random intercept.

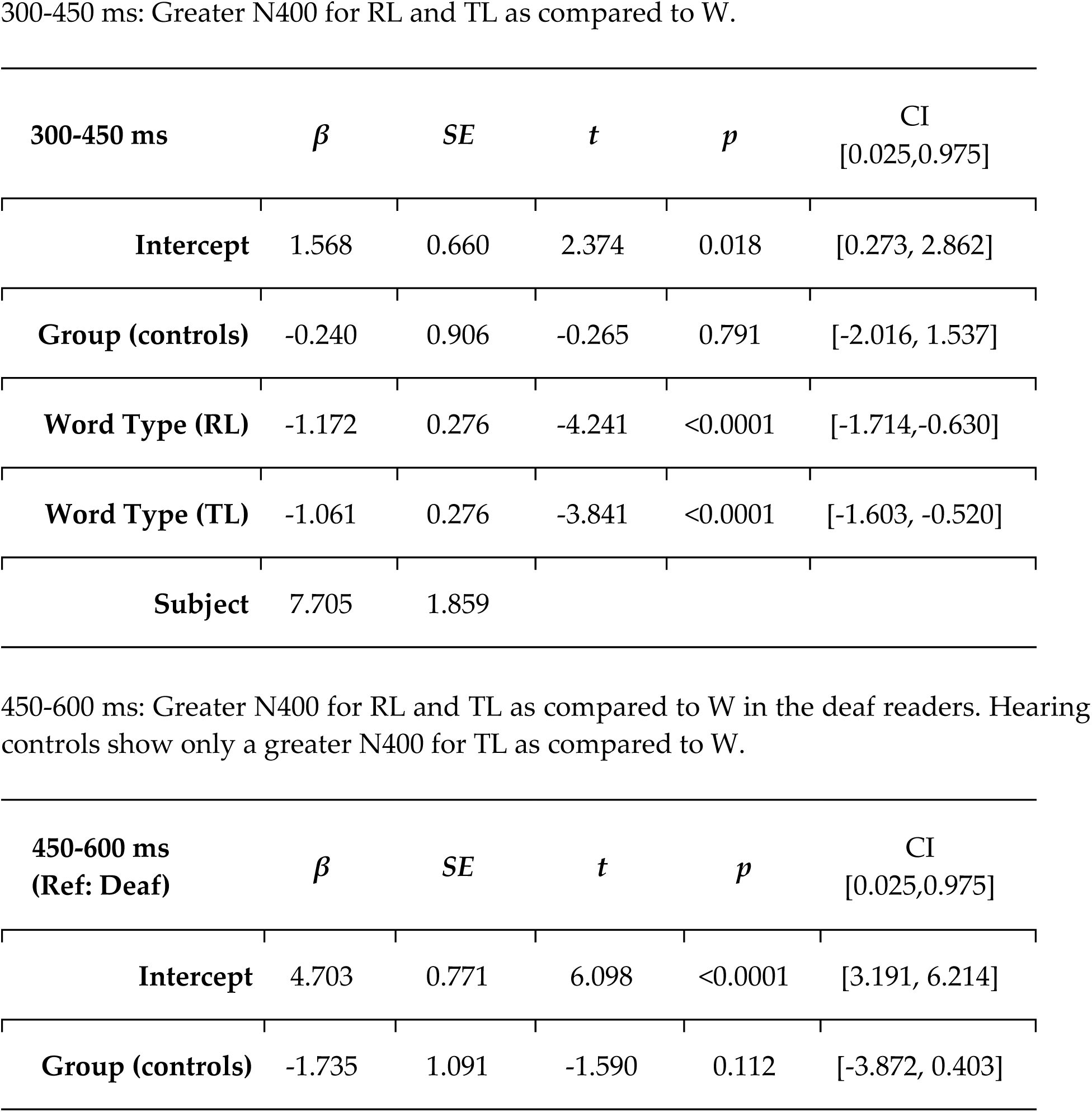
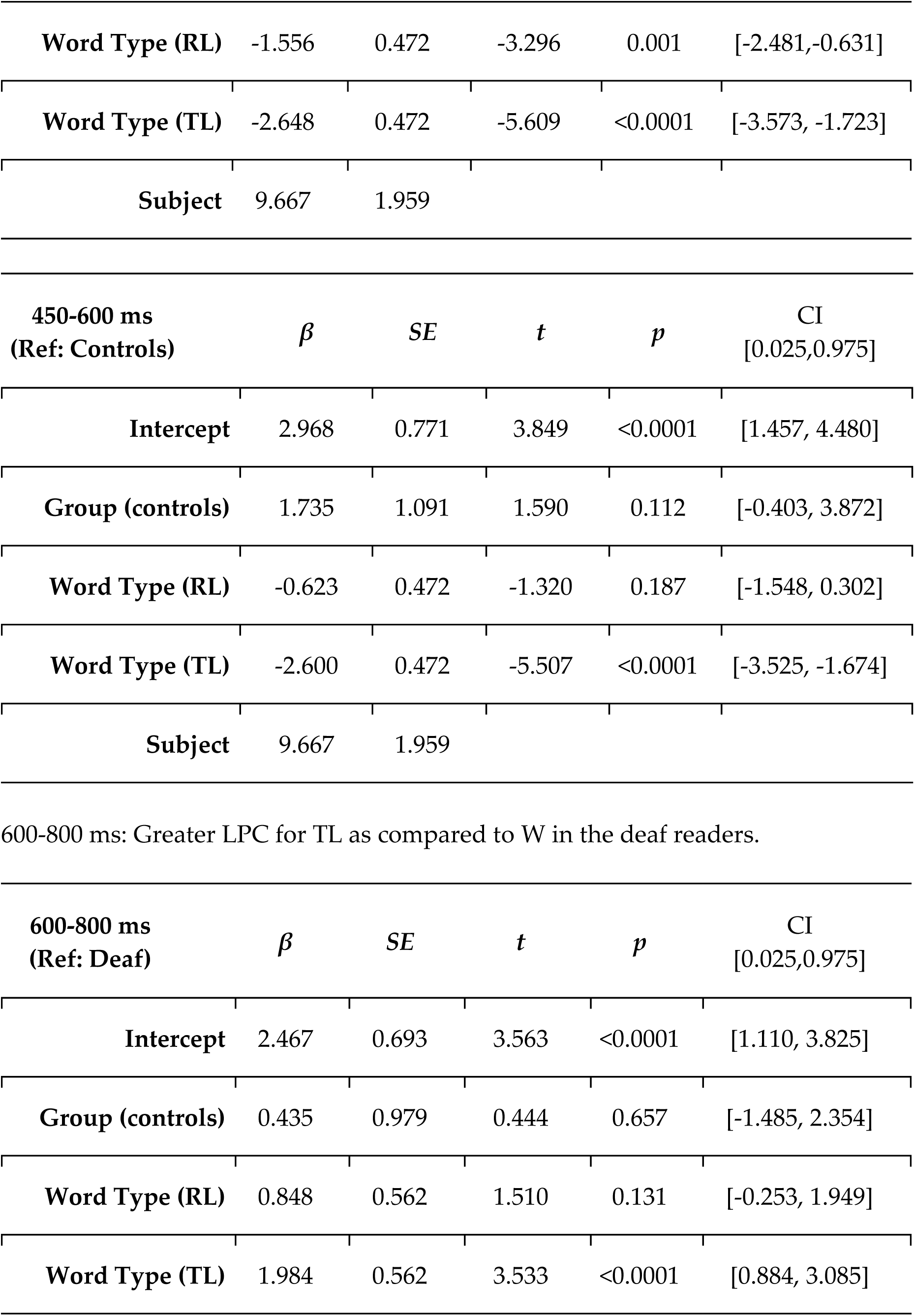
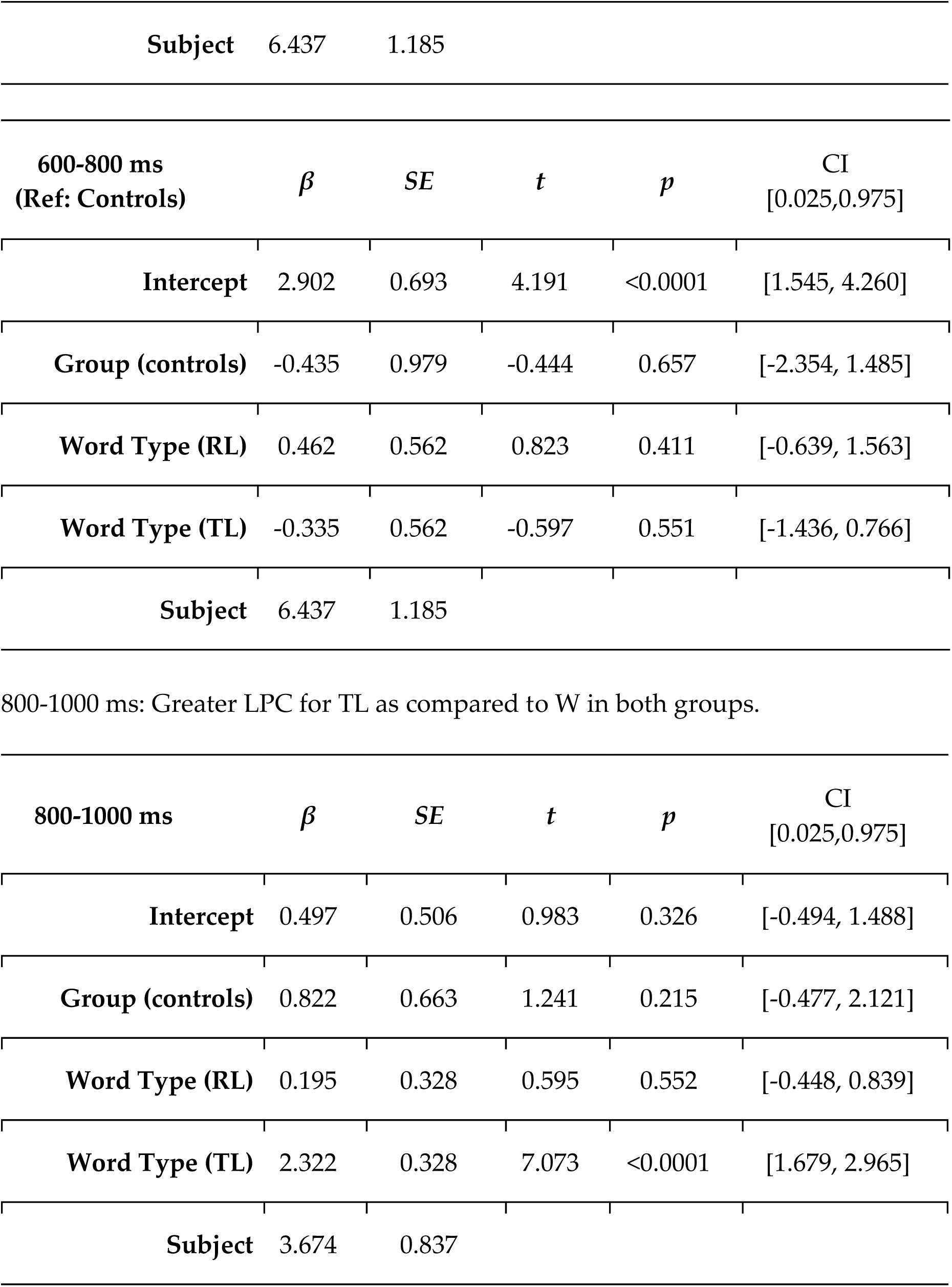

| 300-450 ms | $\beta$ | SE | $t$ | $p$ | CI<br>[0.025,0.975] |
| --- | --- | --- | --- | --- | --- |
| **Intercept** | 1.568 | 0.660 | 2.374 | 0.018 | [0.273, 2.862] |
| **Group (controls)** | -0.240 | 0.906 | -0.265 | 0.791 | [-2.016, 1.537] |
| **Word Type (RL)** | -1.172 | 0.276 | -4.241 | <0.0001 | [-1.714,-0.630] |
| **Word Type (TL)** | -1.061 | 0.276 | -3.841 | <0.0001 | [-1.603, -0.520] |
| **Subject** | 7.705 | 1.859 |  |  |  |

| 450-600 ms<br>(Ref: Deaf) | $\beta$ | SE | $t$ | $p$ | CI<br>[0.025,0.975] |
| --- | --- | --- | --- | --- | --- |
| **Intercept** | 4.703 | 0.771 | 6.098 | <0.0001 | [3.191, 6.214] |
| **Group (controls)** | -1.735 | 1.091 | -1.590 | 0.112 | [-3.872, 0.403] |
| **Word Type (RL)** | -1.556 | 0.472 | -3.296 | 0.001 | [-2.481,-0.631] |
| **Word Type (TL)** | -2.648 | 0.472 | -5.609 | <0.0001 | [-3.573, -1.723] |
| **Subject** | 9.667 | 1.959 |  |  |  |

| **450-600 ms**<br>**(Ref: Controls)** | $\beta$ | *SE* | *t* | *p* | CI<br>[0.025,0.975] |
| --- | --- | --- | --- | --- | --- |
| **Intercept** | 2.968 | 0.771 | 3.849 | <0.0001 | [1.457, 4.480] |
| **Group (controls)** | 1.735 | 1.091 | 1.590 | 0.112 | [-0.403, 3.872] |
| **Word Type (RL)** | -0.623 | 0.472 | -1.320 | 0.187 | [-1.548, 0.302] |
| **Word Type (TL)** | -2.600 | 0.472 | -5.507 | <0.0001 | [-3.525, -1.674] |
| **Subject** | 9.667 | 1.959 |  |  |  |

600-800 ms: Greater LPC for TL as compared to W in the deaf readers.
| **600-800 ms**<br>**(Ref: Deaf)** | $\beta$ | *SE* | *t* | *p* | CI<br>[0.025,0.975] |
| --- | --- | --- | --- | --- | --- |
| **Intercept** | 2.467 | 0.693 | 3.563 | <0.0001 | [1.110, 3.825] |
| **Group (controls)** | 0.435 | 0.979 | 0.444 | 0.657 | [-1.485, 2.354] |
| **Word Type (RL)** | 0.848 | 0.562 | 1.510 | 0.131 | [-0.253, 1.949] |
| **Word Type (TL)** | 1.984 | 0.562 | 3.533 | <0.0001 | [0.884, 3.085] |
| **Subject** | 6.437 | 1.185 |  |  |  |
| **600-800 ms**<br>**(Ref: Controls)** | **$\beta$** | ***SE*** | ***t*** | ***p*** | **CI**<br>**[0.025,0.975]** |
| **Intercept** | 2.902 | 0.693 | 4.191 | <0.0001 | [1.545, 4.260] |
| **Group (controls)** | -0.435 | 0.979 | -0.444 | 0.657 | [-2.354, 1.485] |
| **Word Type (RL)** | 0.462 | 0.562 | 0.823 | 0.411 | [-0.639, 1.563] |
| **Word Type (TL)** | -0.335 | 0.562 | -0.597 | 0.551 | [-1.436, 0.766] |
| **Subject** | 6.437 | 1.185 |  |  |  |
| 800-1000 ms: Greater LPC for TL as compared to W in both groups. |  |  |  |  |  |
| **800-1000 ms** | **$\beta$** | ***SE*** | ***t*** | ***p*** | **CI**<br>**[0.025,0.975]** |
| **Intercept** | 0.497 | 0.506 | 0.983 | 0.326 | [-0.494, 1.488] |
| **Group (controls)** | 0.822 | 0.663 | 1.241 | 0.215 | [-0.477, 2.121] |
| **Word Type (RL)** | 0.195 | 0.328 | 0.595 | 0.552 | [-0.448, 0.839] |
| **Word Type (TL)** | 2.322 | 0.328 | 7.073 | <0.0001 | [1.679, 2.965] |
| **Subject** | 3.674 | 0.837 |  |  |  |

#### Exp 2

LMERs with Prime Type and Group as fixed factors (reference being Same condition) and a by-subject random intercept.

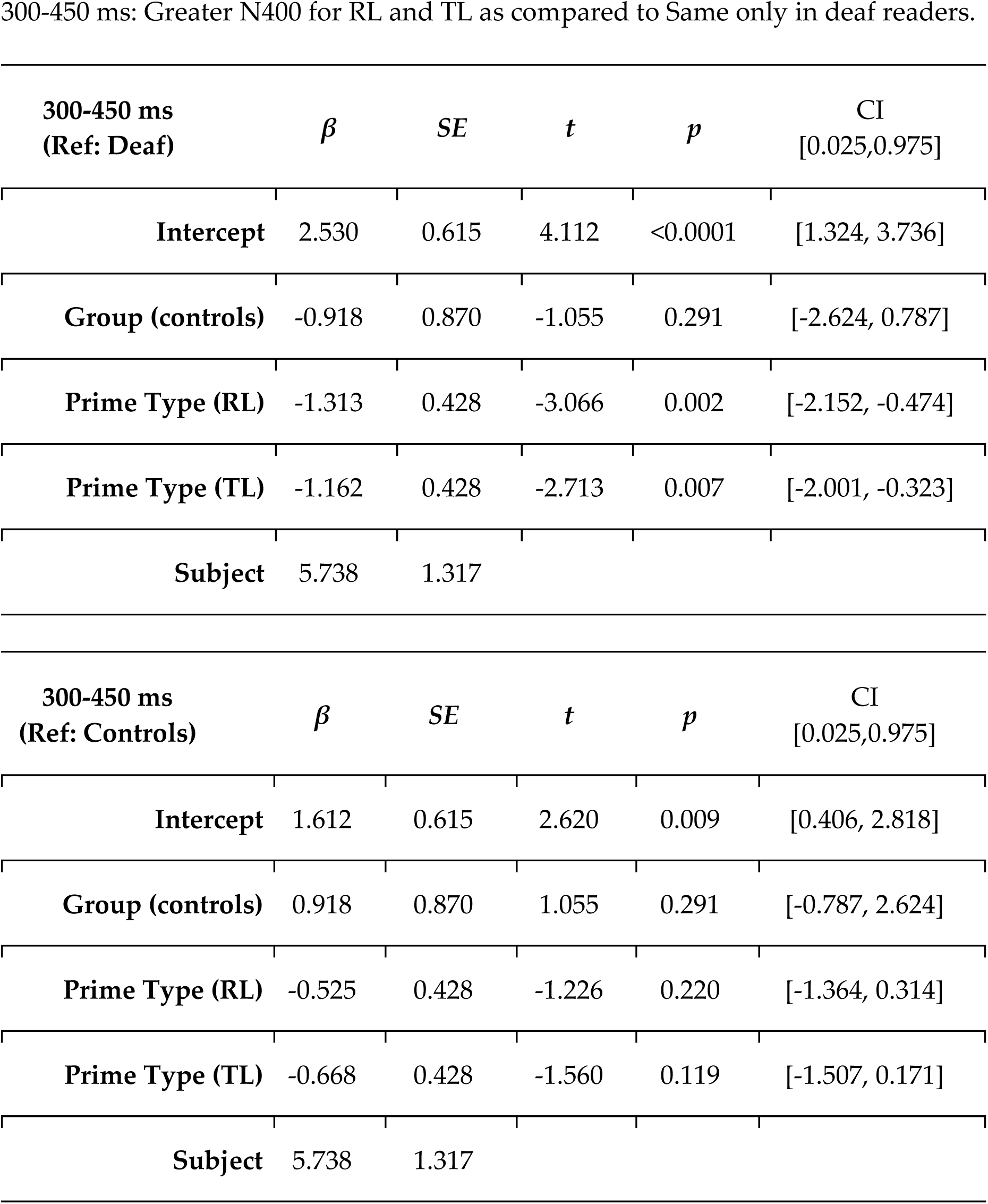
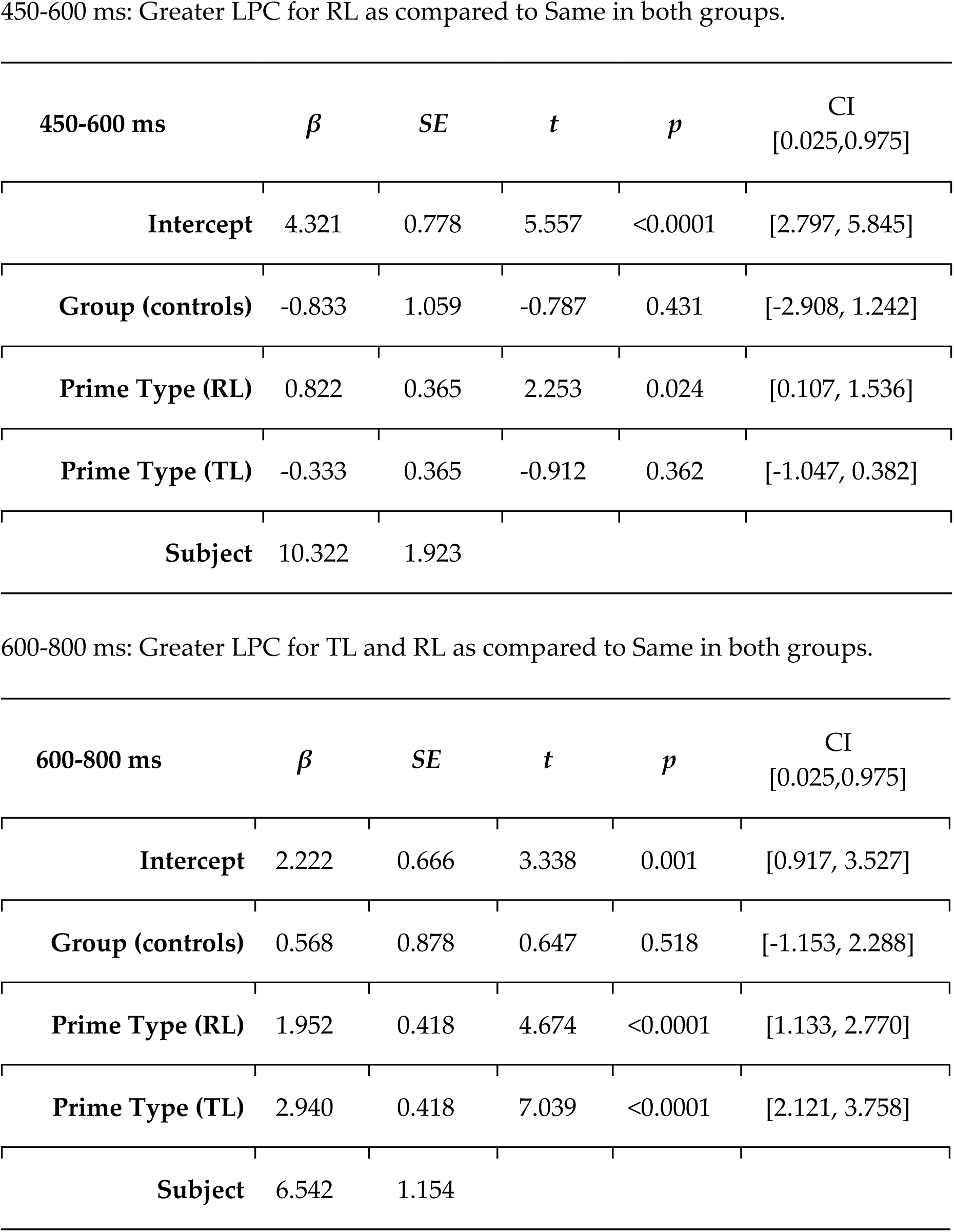
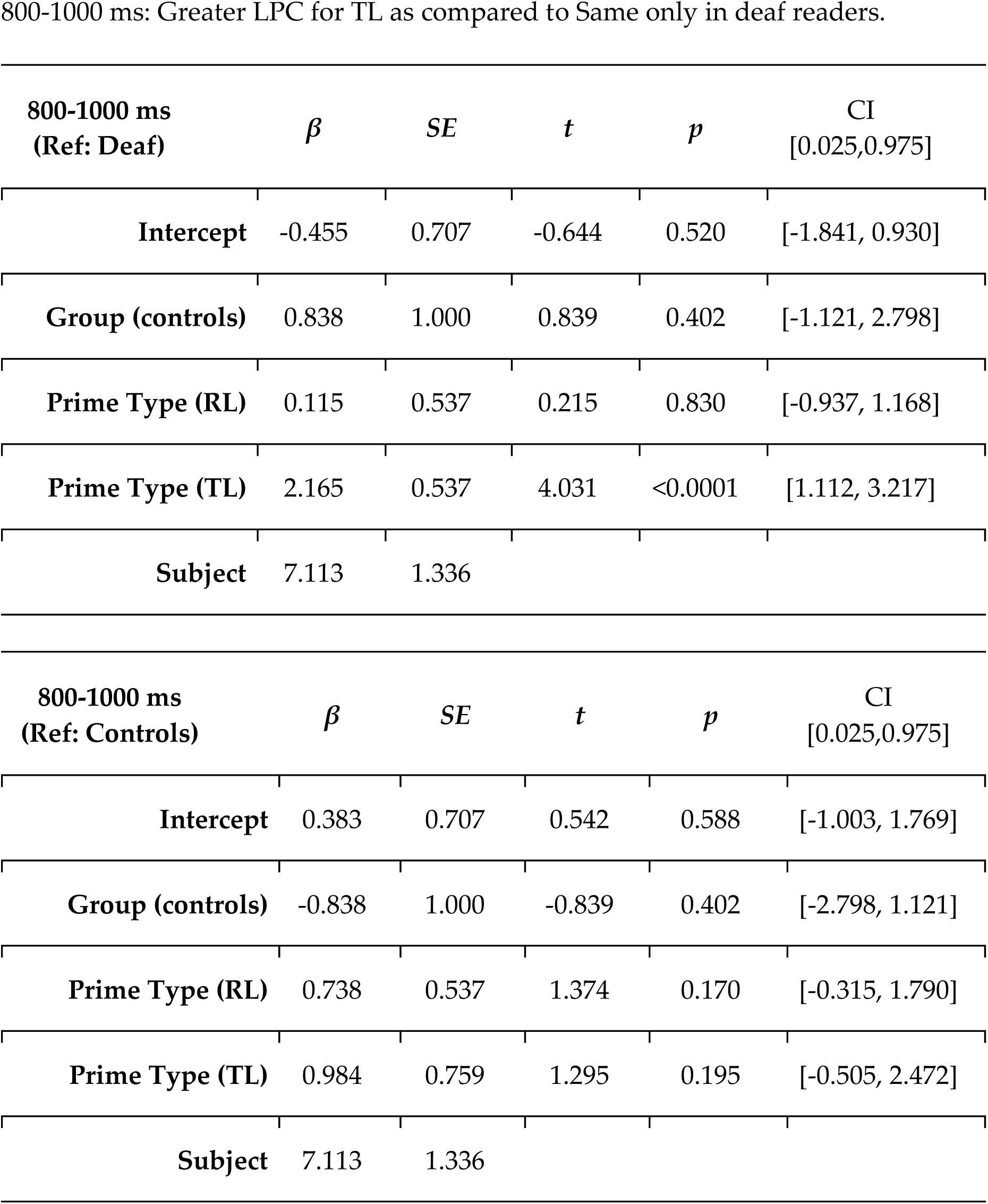

#### Exp 3

LMERs with Prime Type and Group as fixed factors (reference being Same condition) and a by-subject random intercept.

| 300-450 ms<br>(Ref: Deaf) | $\beta$ | SE | $t$ | $p$ | CI<br>[0.025,0.975] |
| --- | --- | --- | --- | --- | --- |
| Intercept | 2.091 | 0.651 | 3.214 | 0.001 | [0.816, 3.366] |
| Group (controls) | -1.027 | 0.920 | -1.116 | 0.264 | [-2.830, 0.776] |
| Prime Type (Diff dl) | -1.155 | 0.384 | -3.004 | 0.003 | [-1.909, -0.402] |
| Prime Type (Diff sl) | -1.290 | 0.384 | -3.355 | 0.001 | [-2.044, -0.537] |
| Subject | 6.987 | 1.729 |  |  |  |
| 300-450 ms<br>(Ref: Controls) | $\beta$ | SE | $t$ | $p$ | CI<br>[0.025,0.975] |
| Intercept | 1.064 | 0.651 | 1.635 | 0.102 | [-0.211, 2.339] |
| Group (controls) | 1.027 | 0.920 | 1.116 | 0.264 | [-0.776, 2.830] |
| Prime Type (Diff dl) | -0.321 | 0.384 | -0.835 | 0.404 | [-1.075, 0.433] |
| Prime Type (Diff sl) | -0.499 | 0.544 | -0.918 | 0.359 | [-1.565, 0.567] |
| Subject | 6.987 | 1.729 |  |  |  |

450-600 ms: No effects of interest.
| 450-600 ms | $\beta$ | SE | $t$ | $p$ | CI<br>[0.025,0.975] |
| --- | --- | --- | --- | --- | --- |
| **Intercept** | 3.974 | 0.774 | 5.137 | <0.0001 | [2.458, 5.490] |
| **Group (controls)** | -0.903 | 1.060 | -0.851 | 0.395 | [-2.981, 1.176] |
| **Prime Type (Diff dl)** | -0.211 | 0.329 | 0.640 | 0.522 | [-0.434, 0.855] |
| **Prime Type (Diff sl)** | -0.515 | 0.329 | -1.567 | 0.117 | [-1.160, 0.129] |
| **Subject** | 10.525 | 2.139 |  |  |  |

600-800 ms: Greater LPC for Diff dl and Diff sl as compared to Same in both groups.
| 600-800 ms | $\beta$ | SE | $t$ | $p$ | CI<br>[0.025,0.975] |
| --- | --- | --- | --- | --- | --- |
| **Intercept** | 1.311 | 0.596 | 2.201 | 0.028 | [0.144, 2.478] |
| **Group (controls)** | -0.153 | 0.813 | -0.188 | 0.851 | [-1.747, 1.441] |
| **Prime Type (Diff dl)** | 0.938 | 0.268 | 3.508 | <0.0001 | [0.414, 1.463] |
| **Prime Type (Diff sl)** | 1.358 | 0.268 | 5.078 | <0.0001 | [0.834, 1.883] |
| **Subject** | 6.140 | 1.547 |  |  |  |

800-1000 ms: Greater LPC for Diff dl as compared to Same in both groups.
| 800-1000 ms | $\beta$ | $SE$ | $t$ | $p$ | CI<br>[0.025,0.975] |
| --- | --- | --- | --- | --- | --- |
| **Intercept** | -0.492 | 0.524 | -0.939 | 0.348 | [-1.520, 0.535] |
| **Group (controls)** | 0.173 | 0.698 | 0.248 | 0.804 | [-1.195, 1.542] |
| **Prime Type (Diff dl)** | 0.277 | 0.305 | 0.906 | 0.365 | [-0.322, 0.875] |
| **Prime Type (Diff sl)** | 0.633 | 0.305 | 2.073 | 0.038 | [0.035, 1.232] |
| **Subject** | 4.254 | 0.999 |  |  |  |

#### Exp 4

LMERs with Prime Type and Group as fixed factors (reference being Same sc condition) and a by-subject random intercept.

| 300-450 ms<br>(Ref: Deaf) | $\beta$ | SE | $t$ | $p$ | CI<br>[0.025,0.975] |
| --- | --- | --- | --- | --- | --- |
| Intercept | 2.090 | 0.595 | 3.515 | <0.0001 | [0.925, 3.256] |
| Group (controls) | -1.021 | 0.841 | -1.214 | 0.225 | [-2.670, 0.627] |
| Prime Type (Diff dc) | -1.100 | 0.318 | -3.453 | 0.001 | [-1.724, -0.475] |
| Prime Type (Diff sc) | -1.084 | 0.318 | -3.403 | 0.001 | [-1.708, -0.459] |
| Prime Type (Same dc) | -0.125 | 0.318 | -0.392 | 0.695 | [-0.749, 0.499] |
| Subject | 6.059 | 1.660 |  |  |  |

| 300-450 ms<br>(Ref: Controls) | $\beta$ | SE | $t$ | $p$ | CI<br>[0.025,0.975] |
| --- | --- | --- | --- | --- | --- |
| Intercept | 1.069 | 0.595 | 1.797 | 0.072 | [-0.097, 2.234] |
| Group (controls) | 1.021 | 0.841 | 1.214 | 0.225 | [-0.627, 2.670] |
| Prime Type (Diff dc) | -0.198 | 0.318 | -0.621 | 0.534 | [-0.822, 0.426] |
| Prime Type (Diff sc) | -0.208 | 0.318 | -0.653 | 0.514 | [-0.832, 0.416] |
| **Prime Type (Same dc)** | 0.137 | 0.318 | 0.431 | 0.666 | [-0.487, 0.761] |
| **Subject** | 6.059 | 1.660 |  |  |  |

| **450-600 ms**<br>**(Ref: Deaf)** | $\beta$ | *SE* | *t* | *p* | CI<br>[0.025,0.975] |
| --- | --- | --- | --- | --- | --- |
| **Intercept** | 3.614 | 0.747 | 4.837 | <0.0001 | [2.150, 5.079] |
| **Group (controls)** | -1.092 | 1.057 | -1.033 | 0.302 | [-3.163, 0.979] |
| **Word Type (Diff dc)** | -1.469 | 0.460 | -3.193 | 0.001 | [-2.371, -0.567] |
| **Word Type (Diff sc)** | -1.172 | 0.460 | -2.547 | 0.011 | [-2.074, -0.270] |
| **Word Type (Same dc)** | 0.378 | 0.460 | 0.822 | 0.411 | [-0.524, 1.280] |
| **Subject** | 9.048 | 1.743 |  |  |  |

| **450-600 ms**<br>**(Ref: Controls)** | $\beta$ | *SE* | *t* | *p* | CI<br>[0.025,0.975] |
| --- | --- | --- | --- | --- | --- |
| **Intercept** | 2.522 | 0.747 | 3.376 | 0.001 | [1.058, 3.987] |
| **Group (controls)** | 1.092 | 1.057 | 1.033 | 0.302 | [-0.979, 3.163] |
| **Prime Type (Diff dc)** | -0.805 | 0.460 | -1.750 | 0.080 | [-1.707, 0.097] |
| **Prime Type (Diff sc)** | -0.724 | 0.460 | -1.572 | 0.116 | [-1.626, 0.178] |
| **Prime Type (Same dc)** | 0.160 | 0.460 | 0.348 | 0.728 | [-0.742, 1.062] |
| **Subject** | 9.048 | 1.743 |  |  |  |

| **600-800 ms** | $\beta$ | *SE* | *t* | *p* | CI<br>[0.025,0.975] |
| --- | --- | --- | --- | --- | --- |
| **Intercept** | 1.981 | 0.574 | 3.452 | 0.001 | [0.856, 3.106] |
| **Group (controls)** | -0.088 | 0.754 | -0.117 | 0.907 | [-1.565, 1.389] |
| **Prime Type (Diff dc)** | 0.522 | 0.348 | 1.500 | 0.134 | [-0.160, 1.205] |
| **Prime Type (Diff sc)** | 1.161 | 0.348 | 3.334 | 0.001 | [0.478, 1.843] |
| **Prime Type (Same dc)** | 1.202 | 0.348 | 3.453 | 0.001 | [0.520, 1.885] |
| **Subject** | 5.073 | 0.963 |  |  |  |

| **800-1000 ms**<br>**(Ref: Same sc)** | $\beta$ | *SE* | *t* | *p* | CI<br>[0.025,0.975] |
| --- | --- | --- | --- | --- | --- |
| **Intercept** | 0.128 | 0.495 | 0.259 | 0.796 | [-0.841, 1.098] |
| **Group (controls)** | 0.040 | 0.653 | 0.061 | 0.951 | [-1.241, 1.321] |
| **Prime Type (Diff dc)** | 0.734 | 0.289 | 2.542 | 0.011 | [0.168, 1.299] |
| **Prime Type (Diff sc)** | 0.993 | 0.289 | 3.440 | 0.001 | [0.427, 1.559] |
| **Prime Type (Same dc)** | 0.846 | 0.289 | 2.932 | 0.003 | [0.280, 1.412] |
| **Subject** | 3.853 | 0.873 |  |  |  |
| **800-1000 ms**<br>**(Ref: Same dc)** | *$\beta$* | *SE* | *t* | *p* | CI<br>[0.025,0.975] |
| **Intercept** | 0.974 | 0.495 | 1.969 | 0.049 | [0.005, 1.944] |
| **Group (controls)** | 0.040 | 0.653 | 0.061 | 0.951 | [-1.241, 1.321] |
| **Prime Type (Diff dc)** | -0.112 | 0.289 | -0.390 | 0.697 | [-0.678, 0.453] |
| **Prime Type (Diff sc)** | 0.147 | 0.289 | 0.509 | 0.611 | [-0.419, 0.713] |
| **Prime Type (Same sc)** | -0.846 | 0.289 | -2.932 | 0.003 | [-1.412, -0.280] |
| **Subject** | 3.853 | 0.873 |  |  |  |

### S3. ERP analysis time locked to the prime onset

The same ANOVAs were applied to EEG data time-locked to the prime onset of Experiment 2, 3 and 4. The time-windows of interest were updated accordingly.

#### Exp 2

##### 600-750 ms

A N400 effect was observed for the non-identical conditions (RL and TL) as compared to the same condition (Prime Type: *F(2,76)*=7.745, *p*<.001; Prime Type x Laterality: *F(4,152)*=4.655, *p*<.01; Prime Type x Anteriority x Laterality: *F(8,304)*=2.768, *p*<.05). A significant interaction Group x Prime Type x Anteriority (*F(4,152)*=2.637, *p*<.05) highlighted that this posterior effect was present only for deaf readers (RL/TL vs Same for deaf readers: *ts*>2, *ps*<.05).

##### 750-900 ms

In both groups the RL condition showed a greater posterior positivity than the other two conditions (Prime Type: *F(2,76)*=4.423, *p*<.05; Prime Type x Anteriority: *F(4,152)*=4.836, *p*<.01; RL differed from TL and Same conditions: *ts*>2, *ps*<.05).

##### 900-1100 ms

In both groups, the non-identical conditions (RL and TL) showed a greater positivity as compared to the same condition (Prime Type: *F(2,76)*=21.819, *p*<.001; Prime Type x Anteriority: *F(4,152)*=4.860, *p*<.01; Prime Type x Laterality: *F(4,152)*=4.078, *p*<.01; Prime Type x Anteriority x Laterality: *F(8,304)*=7.019, *p*<.001).

##### 1100-1300 ms

The TL condition elicited a greater posterior positivity as compared to the other two conditions (Prime Type: *F(2,76)*=9.917, *p*<.001; Prime Type x Anteriority x Laterality: *F(8,304)*=3.870, *p*<.01; TL differed from RL and Same conditions: *ts*>3, *ps*<.01). Critically, this posterior effect was present only in deaf readers (Group x Prime Type x Anteriority: *F(4,152)*=2.517, *p*<.05; deaf effect over central-posterior sites: *ts*>2, *ps*<.05)

#### Exp 3

##### 600-750 ms

A N400 effect was observed for the non-identical conditions (Diff dl and Diff sl) as compared to the same condition (Prime Type: *F(2,76)*=8.553, *p*<.001; Prime Type x Laterality: *F(4,152)*=6.194, *p*<.001; Diff dl/Diff sl vs same: *ts*>3, *ps*<.01). Follow-up analyses of the interaction Group x Prime Type x Laterality showed an asymmetry between groups with a stronger right-lateralized N400 effect only for the deaf readers (deaf: *ts*>3, *ps*<.01).

##### 750-900 ms

None of the contrasts of interest was significant.

##### 900-1100 ms

In both groups, the non-identical conditions (Diff dl and Diff sl) showed a greater positivity as compared to the same condition (Prime Type: *F(2,76)*=8.201, *p*<.001).

##### 1100-1300 ms

None of the contrasts of interest was significant.

#### Exp 4

##### 600-750 ms

Both groups showed an N400 effect for the non-identical conditions (Diff dc and Diff sc) as compared to the same conditions (Same dc and Same sc; Prime Type: *F(1,38)*=17.694, *p*<.001; Prime Type x Anteriority: *F(2,76)*=4.375, *p*<.05; Prime Type x Laterality: *F(2,76)*=4.683, *p*<.05). This effect was stronger and wider distributed for deaf readers (Group x Prime Type x Anteriority x Laterality: *F(4,152)*=3.411, *p*<.05; the deaf group showed a difference over anterior, central and posterior sites: *ts*>3, *ps*<.01, while the hearing controls showed a difference only over posterior sites: *ts*>3, *p*<.05).

##### 750-900 ms

An N400 effect persisted over central-posterior sensors (Prime Type: *F(1,38)*=15.006, *p*<.001; PrimeType x Anteriority: *F(2,76)*=11.791, *p*<.001).

##### 900-1100 ms

Both groups showed a greater positivity for the non-identical conditions (Diff dc and Diff sc) as compared to the same conditions (Prime Type x Anteriority x Laterality: *F(4,152)*=2.601, *p*<.05). This effect was wider distributed for the same case strings (Prime Type x Case: *F(1,38)*=9.972, *p*<.01; Prime Type x Case x Anteriority: *F(2,76)*=11.576, *p*<.001; same cases showed a difference over central posterior sensors, *ts*>2, *ps*<.05, while different cases showed a marginal difference only over the cluster of posterior sensors, *t*>2, *p*=.09).

##### 1100-1300 ms

The central posterior positive effect persisted only for the same case string (Prime Type x Case: *F(1,38)*=5.529, *p*<.05; Prime Type x Case x Anteriority: *F(2,76)*=5.354, *p*<.05; Prime Type x Case x Laterality x Anteriority: *F(4,152)*=4.495, *p*<.01; central and posterior sensors, *ts*>2, *ps*<.05).

